# Microglia/macrophage-specific deletion of TLR-4 protects against neural effects of diet-induced obesity

**DOI:** 10.1101/2024.02.13.580189

**Authors:** Jiahui Liu, Ali Zaidi, Christian J. Pike

## Abstract

Obesity is associated with numerous adverse neural effects, including reduced neurogenesis, cognitive impairment, and increased risks for developing Alzheimer’s disease (AD) and vascular dementia. Obesity is also characterized by chronic, low-grade inflammation that is implicated in mediating negative consequences body-wide. Toll-like receptor 4 (TLR4) signaling from peripheral macrophages is implicated as an essential regulator of the systemic inflammatory effects of obesity. In the brain, obesity drives chronic neuroinflammation that involves microglial activation, however the contributions of microglia-derived TLR4 signaling to the consequences of obesity are poorly understood. To investigate this issue, we first generated mice that carry an inducible, microglia/macrophage-specific deletion of TLR4 that yields long-term TLR4 knockout only in brain indicating microglial specificity. Next, we analyzed the effects of microglial TLR4 deletion on systemic and neural effects of a 16-week of exposure to control versus obesogenic high-fat diets. In male mice, TLR4 deletion generally yielded limited effects on diet-induced systemic metabolic dysfunction but significantly reduced neuroinflammation and impairments in neurogenesis and cognitive performance. In female mice maintained on obesogenic diet, TLR4 deletion partially protected against weight gain, adiposity, and metabolic impairments. Compared to males, females showed milder diet-induced neural consequences, against which TLR4 deletion was protective. Collectively, these findings demonstrate a central role of microglial TLR4 signaling in mediating the neural effects of obesogenic diet and highlight sexual dimorphic responses to both diet and TLR4.

## Introduction

Obesity is a global health concern that is associated with increased risks for impaired functions and development of disease in tissues throughout the body, including metabolic syndrome^1^, diabetes^2^, cardiovascular disease^3^, and certain cancers^4^. In the brain, obesity is associated with decreased hippocampal volume^5^, impaired cognition, increased risk of dementia^6–8^ and other deleterious outcomes. The mechanisms by which obesity adversely affects tissues are many, including evidence that obesity accelerates pathological processes associated with aging ^9,10^. One of the mechanisms contributing to the adverse effects of obesity is inflammation, particularly chronic, low-grade inflammation that affects numerous tissues, including the brain^11^.

Toll-like receptor 4 (TLR4) signaling is crucial in mediating inflammatory events^12,13^. In humans, the metabolic disorders insulin resistance and obesity have been associated with increased TLR4 levels in many tissues, including peripheral blood mononuclear cells^14,15^, adipose tissue^15,16^, and muscle^17^. Importantly, a recent study showed that TLR4 was required for saturated fatty acid (SFA)-induced inflammation through cellular metabolism reprogramming^18^. A central role of TLR4 in mediating obesity-related outcomes is supported by animal studies showing that TLR4 loss-of-function mutation, knockdown or pharmacological inhibition, protected mice systemically from diet-induced inflammation and insulin resistance^19–24^. TLR4 signaling is also linked with adverse effects of obesity in brain, as protection from neural effects of obesogenic diet have been reported in mice treated systemically with a pharmacologic TLR4 inhibitor and in mice with a loss-of-function TLR4 mutation^25,26^. However, the specific contributions of neural versus peripheral TLR4 signaling in driving neural outcomes of obesity remain incompletely understood.

In the central nervous system (CNS), TLR4 expression is predominantly observed in microglia under both basal conditions and following exposure to obesogenic high-fat diet (HFD) ^13,27,28^. As with other tissues, activation of TLR4 signaling in brain is associated with inflammation^29^. Neuroinflammation is widely considered as a key mediator of obesity-induced neural dysfunction^30–32^. At the cellular level, obesity drives neuroinflammation in part by increasing microglial production of pro-inflammatory cytokines in hypothalamus, hippocampus, and other brain regions^33–35^. Microglia proliferation and activation, as well as altered morphology were observed after HFD exposure^25,36^. Studies targeting neuroinflammation by depleting microglia or inhibiting microglial activation demonstrated that microglia modulate food intake during HFD feeding and contribute to diet-induced obesity (DIO)^37–39^.

Collectively, the existing literature suggests that microglia-dependent TLR4 signaling may be a primary mechanism underlying deleterious effects of obesity in the brain, however this has not yet been directly tested. To investigate the role of microglial TLR4 signaling in obesity-induced neural impairments, we generated a novel transgenic mouse model with inducible deletion of TLR4 specifically within cells expressing CX3CR1, which are essentially limited to microglia in CNS and macrophages peripherally^40–42^. This approach has been successfully used to achieve long-term deletion of target genes primarily within microglia due to the rapid turnover of most peripheral macrophage populations^43^. In this study, we compared metabolic, molecular, and behavioral outcomes following a 16-week exposure to control diet or high-fat diet in male and female mice carrying the microglial TLR4 deletion versus a transgenic control line.

## Methods

### Animals and Treatment

Microglia specific TLR4 knockout mice were generated by crossing *TLR4*^flx/flx^ mice (Jackson Labs, Stock No: 024872) with CX3CR1^CreER^ mice (Jackson Labs, Stock No: 020940), which express CreER recombinase induced by tamoxifen administration under the control of CX3CR1 promoter. *TLR4* flox-homozygous and Cre-positive mice (*TLR4*^flx/flx^, CX3CR1^CreER +/−^) are referred to as the TLR4-MKO model. Their littermates, which are Cre-negative (*TLR4*^flx/flx^, CX3CR1^CreER −/−^) served as the control group (Ctl). At 3 months of age, male and female mice were treated with tamoxifen (i.p., 150 mg/kg, Sigma Aldrich; solubilized in safflower oil) at two time points 48 hr apart, a regimen previously reported to induce gene deletion^43^. Two weeks following the last treatment, mice within each line were randomly assigned into two nutrient-matched dietary groups: control diet (10% fat; #D12450J, Research Diets; CD) or high fat diet (60% fat; #D12492, Research Diets; HFD). Animals were maintained on experimental diets for 16 weeks and were housed under 12h light/dark cycle with lights on at 0600h with *ad libitum* access to food and water.

At the conclusion of the dietary treatment, animals were euthanized via carbon dioxide exposure following a 16h overnight fasting. Blood samples were collected by cardiac puncture into EDTA-coated tubes. Aliquots of whole blood and plasma were frozen at −80 °C. Animals were perfused with ice-cold phosphate-buffered saline (PBS) for 15 min. Brain was rapidly removed and either dissected into cerebral cortex, hippocampus and hypothalamus and immediately frozen at −80°C, or fixed in 4% paraformaldehyde in 0.1 M Sorenson’s phosphate buffer for 72 h at 4°C. Visceral fat pad was also harvested and stored at −80 °C.

### Metabolic Measurements

Body weight and food intake were recorded on a weekly basis during the 16-week diet period. Body composition (lean mass, fat mass and fluid) was determined using an NMR analyzer (Bruker LF90 Minispec, Bruker Optics).

### Glucose Measurement

Fasting glucose was measured at week 0 and every 4 weeks afterwards of the 16-week diet period following an overnight fast (16h). Blood was collected from the lateral tail vein and immediately measured for glucose level using a Precision Xtra Glucose Monitor (Abbott). At week 12, a glucose tolerance test (GTT) was performed immediately after baseline fasting glucose level was taken. Mice were administered a glucose bolus (2 g/kg, D-glucose) via i.p. injection and blood glucose levels were recorded 15, 30, 60, and 120 min thereafter.

### Behavior

For all behavioral tests, mice were brought into the behavior room and allowed to acclimate for 30 min prior to testing. After each trial, animals were returned to their home cages. The testing arena was disinfected with 70% ethanol and air-dried before the next trial. The elevated plus maze test was scored live. All other tests were recorded by a video camera mounted above the maze and analyzed using Noldus EthoVision XT software (Version 14). Both camera and software were calibrated according to the manufacturer’s instructions. All behavioral tests were performed during the hours of 0800 - 1600.

Open field test was performed 1 week following the last tamoxifen treatment. After 30 min acclimation, mice were placed in the middle of the arena (40 cm × 40 cm) and allowed to move freely for 5 min. The arena was equally divided into 16 squares and the middle 4 squares were identified as center area. Total distance moved, total time the animals are mobile, and time spent in center were recorded. Average velocity was calculated as total distance moved divided by total time mobile.

Elevated plus maze was performed one day after the open field test. The maze was elevated 40 cm above the floor and consisted of a center area (6 cm × 6 cm), two closed arms (30 cm × 6 cm) and two open arms (30 cm × 6 cm). Briefly, mice were placed in the center platform of the maze facing a closed arm and allowed to move freely for 5 min. An arm entry or exit was defined as both front paws placed into or outside of the arm. Arm choices and time spent in open arms were recorded for each animal.

Barnes maze was performed at week 14 of dietary treatment using modifications of a previously described protocol ^44^. The maze was 90 cm above the floor and had a white circular platform (91.5 cm in diameter) with 20 circular holes (5 cm in diameter) evenly spaced around the border. The maze was walled on all four sides with black curtains. One unique visual cue was placed on each wall at the level of the platform. On day 1, animals were habituated to the maze. Mice were placed in the center of the maze and allowed to move freely for 3 min under red light in an otherwise dark room. All holes were closed during the habituation trial. From days 2 - 5, each animal was given three training trials per day with a 15 min inter-trial interval. One cuboid escape box (11 cm L x 5 cm W x 5 cm H) was hidden beneath the maze. Three decoy boxes (5 cm L x 5 cm W x 2.5 cm H) were also placed beneath the maze to prevent visual confirmation of the location of the escape box. All other holes were closed during the training trials. The location of the escape and decoy boxes varied among animals but were kept constant for a given mouse throughout all trials. A bright light was placed directly above the maze, and a buzzer located below the maze was turned on during the task to stimulate entry into the escape box. During each training trial, mice were placed in an opaque cylinder in the middle of the maze. After 10 s delay the cylinder was lifted, and the animals were allowed to move freely for up to 3 min to locate and enter the escape box. If the animals did not find the escape box within 3 min, they were gently hand-guided into the box. Upon entry into the escape box either by successful escape or guidance, the light and buzzer were turned off and mice remained in the escape box for 1 min before being returned to their home cages. In each trial, the latency for their full body to enter the escape box was recorded. Average latency of three training trials performed on the same day was calculated for each animal.

Forty-eight hours after the last training trial (day 7), mice were tested on a probe trial in which the escape box was switched to a decoy box. The location of other three decoy boxes remained unchanged from training trials. The animals were placed in the cylinder for 10 s. Then the cylinder was lifted and the mice were allowed to freely explore the maze for 3 min. Correct holes were considered as the target hole where the escape box was previously located, as well as one adjacent hole on each side. Latency to reach the target hole (nose poke) for the first time, and number of errors (nose pokes into incorrect holes) were recorded. Two animals, one male and one female from the TLR4-MKO group that were fed on CD, did not show successful escapes over the four days of training. Consequently, these animals were excluded from all analyses due to unresponsiveness to the behavioral task. The exclusion did not affect the statistical results.

Novel object placement and recognition tests were performed two days after the probe trial in Barnes maze test, with adaptations of a previously described protocol ^45,46^. The same arena as the open field test was used. To reduce potential anxiety associated with exposure to test objects, a small plastic toy block was placed in the home cage of each animal one day prior to the habituation trial and remained there until all test trials were completed. Two sets of objects (metal padlock, approximately 4 cm L x 4.5 cm W x 2.5 cm H; and small stapler, approximately 5 cm L x 3 cm W x 4.5 cm H) were used in sampling and test trials. On days 1 and 2, each animal was given one habituation trial per day for 2 consecutive days. During habituation, mice were placed in the middle of the empty open field arena and allowed to freely explore the maze for 5 min. On day 3, mice were given one habituation trial, one sampling trial and one test trial. Twenty-four hours after the second habituation trial (day 3), animals were placed in the middle of the empty arena to freely explore for 2 min. Next, the arena was immediately set up with two identical objects placed in two adjacent corners (5 cm away from either wall). Mice were placed back in the arena with their heads positioned opposite the objects. The trial lasted for 20 min or until a criterion that the combined exploration time of 30 s of both objects was met. Mice were considered to be exploring an object when showing investigative behavior while their nose is within 1.5 cm of the object. *Novel object placement*: Four hours after the sampling trial, animals were placed in the arena with one of the identical objects moved to the diagonally opposite corner. The animals were given 20 min or until they had explored the two objects for a combined total of 30 sec. *Novel object recognition*: On day 4 (24 h after the sampling trial), animals were placed in the arena with one of the identical objects replaced by a novel object. Animals were given 20 min or until the criterion of 30 s of total exploration time of both objects was achieved. During sampling and test trials, the exploration time of each object was recorded. The location and nature of the novel object were randomized and balanced across all animals. In pilot studies, the animals did not show significance preference for either of the two objects (data not shown).

### Blood and Plasma Measurements

Glycated hemoglobin (Hba1c) levels were measured in whole blood using a commercially available kit (80310; Crystal Chem). Fasting leptin and insulin were measured in plasma by ELISA kits (EZML-82K, MilliporeSigma; EZRMI-13K, MilliporeSigma). Homeostatic model for insulin resistance (HOMA-IR) was calculated using the formula: [ fasting blood glucose (mg/dL) × fasting plasma insulin (mU/L) / 405]. Plasma cytokine levels were measured using Meso Scale Discovery (MSD) V-Plex Proinflammatory Panel 1 (mouse) cytokine assay and analyzed on a QuickPlex SQ 120 instrument (MSD). Levels of IL-1β, IL-4 and IL-12p70 were below fit curve range in over half of the samples and therefore were excluded. All measurements were performed according to the manufacturer’s protocol. All standards and samples were measured in duplicate.

### Immunohistochemistry

Fixed brains were transferred into 20% sucrose in PBS for 2 days until they sank down to the bottom. Brains were then exhaustively sectioned in the coronal plane at 20 μm using a cryostat (Leica Biosystems). Sections were stored in PBS with 0.03% sodium azide at 4°C until immunohistochemistry was performed. Every fourth section from approximately bregma −1.40 mm to −2.00 mm were immunostained with ionized calcium binding adaptor molecule 1 (Iba1; 1:2000; FUJIFILM Wako) and glial fibrillary acidic protein (GFAP; 1:1000; Dako); 4-6 sections per brain. Every fourth section from approximately bregma −2.40 mm to −2.90 mm was immunostained with doublecortin (DCX; 1:2500; Santa Cruz); 4-6 sections per brain. For Iba1 staining, brain sections were first incubated with 10 mM EDTA (pH 6) at 95°C for 10 min. For DCX staining, brain sections were pre-treated with 95% formic acid for 5 min at room temperature. No antigen retrieval pretreatment was performed for GFAP staining. Next, sections were rinsed in Tris-buffered saline (TBS) and treated with an endogenous peroxidase blocking solution for 10 min. Sections were then rinsed in TBS with 0.2% Triton-X before being blocked for 30 min in corresponding blocking solution. The blocking solution consisted of TBS with 2% bovine serum albumin (BSA) for Iba1, 2% BSA and 2% normal goat serum for GFAP, and 5% normal horse serum for DCX. Sections were incubated overnight at 4°C with primary antibodies diluted in their respective block solutions. On the following day, sections were rinsed in TBS with 0.1% Triton-X and incubated in appropriate biotinylated secondary antibody diluted in the blocking solution for 1 h. After rinsing in TBS with 0.1% Triton-X, sections were incubated in an avidin-biotin complex (Vectastain ABC Elite kit, Vector Laboratories) for 1 h and immunoreactivity was visualized using diaminobenzidine tetrahydrochloride (Vector Laboratories).

Microglia morphology was analyzed as a measure of activation phenotype as previously described^47^. Briefly, brightfield images of Iba-1 immunostained sections containing the hippocampus CA1 (2 fields/section, 4 sections/animal) or arcuate nucleus (ARC) of the hypothalamus (1 field/section, 6 sections/animal) were collected using a Keyence BZ-X710 microscope. Z series stacks (at least 10μm) with 0.7μm interval were acquired under 40x objective and processed with maximum contrast projection using BZ-X Analyzer software (Version 1.3.1.1, Keyence). Images were imported into ImageJ (Version 1.53m) and regions of interest (ROIs) of the CA1 and ARC were manually outlined. Microglia process length and endpoints were analyzed using the AnalyzeSkeleton (2D/3D) plugin^48^. Microglia cell somas (5 cells/field, 40 cells/animal) were manually outlined, and their size and roundness were determined using ImageJ (Version 1.53m).

To quantify GFAP immunoreactivity, brightfield images containing the hippocampus (20-30 fields/section, 4 sections/animal) and ARC (1 field/section, 6 sections/animal) were captured with Keyence BZ-X710 microscope under 20x objective. For hippocampus, overlapping images from different fields were merged (uncompressed) using BZ-X Analyzer software (Version 1.3.1.1, Keyence). ROIs of the entire CA1 and ARC were manually outlined and images were converted to greyscale and thresholded using ImageJ (Version 1.53m). GFAP immunohistochemical load was calculated as the percentage of the positively immunolabeled pixels over the total area of ROI.

DCX+ cells in the sub-granular zone and granule cell layer of the dentate gyrus (4 sections/animal) were counted live using Olympus BX50 microscope under 100× objective. DCX+ cells were classified into subtypes based on morphology as described in previous studies^49–51^: type 1 - no or one short (shorter than the diameter of the cell body) process; type 2 - one process longer than type 1 but only reaching within the granule cell layer; or type 3 - one long process or multiple processes that branch into the molecular layer. The percentages of type 1, 2, and 3 cells were calculated for each animal.

### Microglia and monocyte isolation

Microglia and peripheral monocytes were collected from 6-7 month old male and female mice 3 weeks after last tamoxifen treatment.

Animals were euthanized via carbon dioxide exposure and perfused with ice-cold PBS for 5 min. Brains were quickly removed and placed into Hanks’ balanced salt solution without Ca^2+^, Mg^2+^ (HBSS w/o) on ice. Whole brain from each mouse was cut into small pieces using a sterile scalpel blade. After centrifugation at 300 × *g* for 2 min, brain tissues were homogenized using a neural tissue dissociation kit (MiltenyiBiotec) following the manufacturer’s instruction. The resultant cell suspension was passed through a cell strainer (70 µm; MiltenyiBiotec) and then centrifuged at 300 x *g* for 10 minutes at 4°C. Pelleted cells were resuspended into 30% Percoll (GE Healthcare) and centrifuged at 700 × *g* for 15 minutes at room temperature. After centrifugation, an upper myelin layer was carefully removed. Cells were resuspended in MACS buffer (PBS with 0.5% BSA and 2 mM EDTA, pH 7.2) and incubated with CD11b microbeads (MiltenyiBiotec) for 15 minutes at 4°C. After washing with MACS buffer, cells were loaded onto columns (MiltenyiBiotec) and separated using a magnetic separator (MiltenyiBiotec). Microglia (CD11b+) and all other cells (CD11b-) were collected and centrifuged at 300 x *g* for 10 min. Cell pellets were stored at −80°C.

Peripheral monocytes were isolated using adaptation of a described protocol^52^. Briefly, for each mouse blood (0.8-1ml) was collected by cardiac puncture into EDTA-coated tubes. Blood was then diluted with red blood cell lysis solution (MiltenyiBiotec) and incubated 10 minutes at room temperature. After centrifuging at 300 × *g* for 10minutes, cells were resuspended in Roswell Park Memorial Institute (RPMI) medium and carefully layered on top of Ficoll (GE Healthcare). The upper layer of plasma was carefully removed, and the middle layer containing mononuclear cells was collected. Subsequently, the cells were washed and suspended in HBSS before undergoing centrifugation at 400 × *g* for 30 minutes. Following another round of centrifugation at 200 × g for 5 minutes, the cells were resuspended in RPMI medium and transferred to cell culture plates. After 30 min incubation in a 5% CO_2_ container at 37°C, nonadherent cells were discarded and adherent cells (monocyte fraction) were collected after washing with PBS.

### RNA isolation and quantitative PCR

RNAs from isolated monocytes, microglia and other brain cells were extracted using the quick-RNA miniprep kit (Zymo Research) according to manufacturer’s instructions. For adipose and brain tissues, RNA was extracted using TRIzol reagent (Invitrogen) following the manufacturer’s protocol, except an extra step of centrifuging at 12000 × *g* for 10 min was performed after homogenization in TRIzol for adipose tissue to remove excess lipid. Purified RNA (500 ng) was reverse-transcribed using the iScript cDNA synthesis system (Bio-Rad). The resulting cDNA was used for real-time quantitative PCR using SsoAdvanced Universal SYBR Green Supermix (Bio-Rad) and the Bio-Rad CFX Connect Thermocycler. All samples were tested in duplicates. Expression level of each probed gene was compared with averaged Ct value of succinate dehydrogenase complex, subunit A (SDHA) and hypoxanthine guanine phosphoribosyl transferase (HPRT) in the adipose tissue, β-actin and phosphoglycerate kinase 1 (pgk1) in the brain tissue, and β-actin in the isolated microglia and monocytes. Relative quantification of mRNA was determined using the ΔΔ-CT method and normalized to Ctl mice or male Ctl mice fed on CD. Data was presented as log2 fold change and statistical test was run using the ΔCT values. Primer pair sequences of all target genes are listed in Table 1.

**Table 1.**
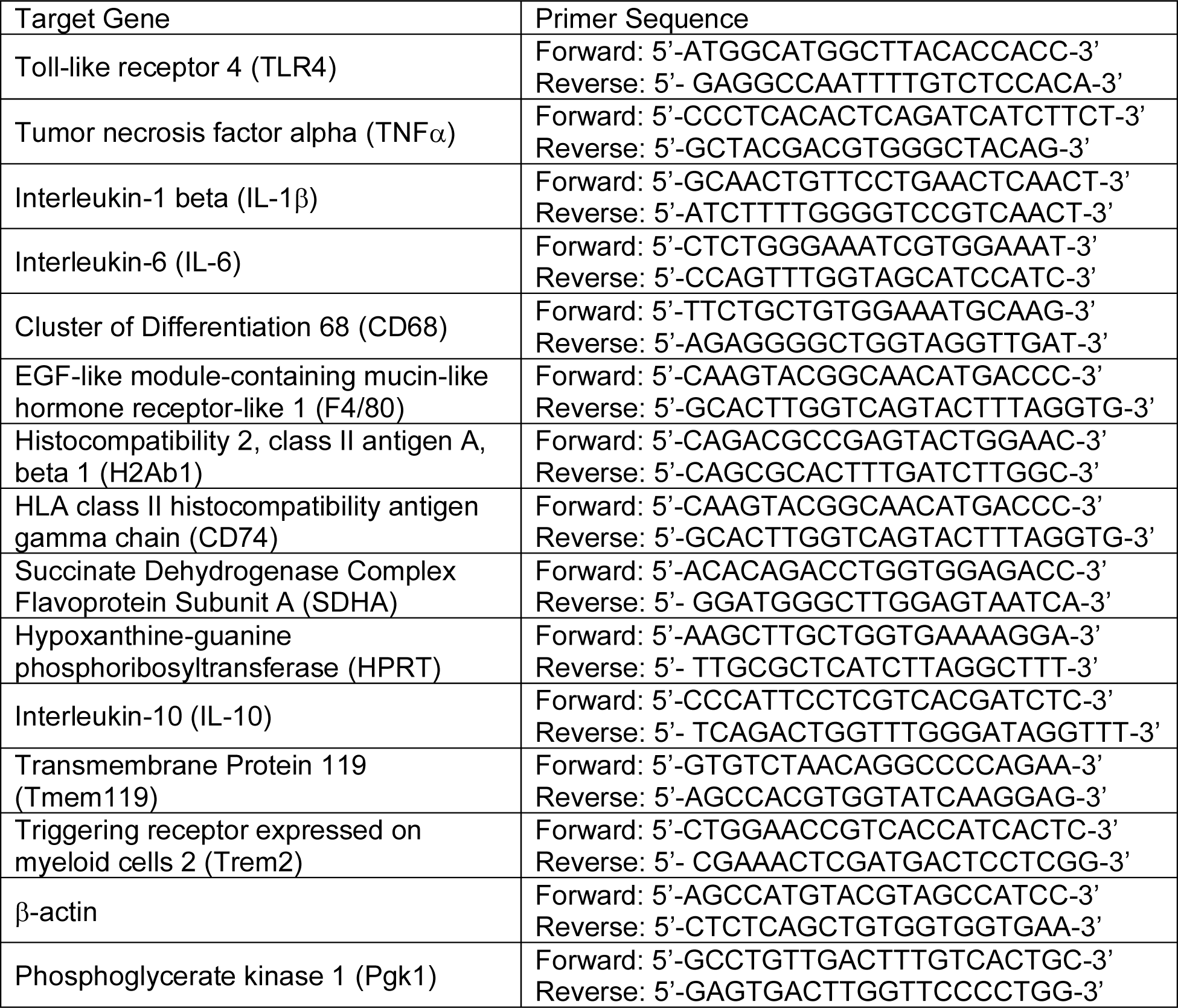
Target genes and their respective PCR primer sequences.

### 2.9 Statistical analyses

Data were analyzed using GraphPad Prism (Version 9) and SPSS (Version 28). RNA expression in isolated microglia and monocyte, open field test and elevated plus maze were analyzed by unpaired two-tailed *t*-test with Welch’s correction. Novel object placement and recognition tests were analyzed using one-sample *t*-test to determine whether the time spent with the moved or novel object differed from chance level (15s)^45,53^. Three-way repeated ANOVAs were performed on body weights, fasting glucose, glucose tolerance and escape latency in the Barnes maze test with repeated measures across time points and genotype and diet as a between-subjects factors. All other data were analyzed with two-way ANOVAs with genotype and diet as between-subjects factors. For all ANOVAs, Greenhouse-Geisser correction was used and significant effects were further analyzed with planned comparisons between groups of interest using the Bonferroni or Tukey *post hoc* testing. All cytokine levels in plasma except for TNFα showed a highly non-normal distribution by Kolmogorov-Smirnov and Shapiro-Wilks tests (p < 0.001), and skewness was either greater than 1 or less than −1. Thus, data were log10 transformed before analysis. All datasets are expressed as mean ± SEM or from minimum to maximum with quantiles. The threshold for statistical significance was set at *p* < 0.05.

## 3. Results

### 3.1 Microglia TLR4 deletion improved HFD-induced insulin resistance without affecting adiposity in male mice

To assess the role of microglial TLR4 signaling in mediating the effects of obesogenic diet, we created a transgenic mouse model in which we induced TLR4 deletion at adult age largely within microglia by taking advantage of the different turnover rates in different CX3CR1+ cell populations. It has been demonstrated that CX3CR1 is exclusively expressed by microglia in the adult brain^54^. Due to their longevity and mostly self-proliferation during microgliosis^55^, it has been reported that gene recombination in microglia remains stable over several months after tamoxifen treatment in the CX3CR1^CreER^ model^43^. During adulthood CX3CR1 is also expressed in circulating blood monocytes, as well as other subsets of peripheral mononuclear phagocytes, including tissue macrophages and dendritic cells^56,57^. However, these cells generally are short-lived and replaced by bone marrow-derived progenitors^40^, yielding cells expressing unarranged TLR4. To validate TLR4 gene deletion, we isolated microglia and blood mononuclear myeloid cells from TLR4-MKO and Ctl mice 3 weeks after completing tamoxifen injections. After tamoxifen treatment, mRNA level of TLR4 in TLR4-MKO mice was reduced in microglia but comparable to Ctl mice in other brain cells and peripheral monocytes (Figure S1). Acute tamoxifen treatment has been reported to result in short-term changes in body composition without affecting body weight, along with inducing enduring changes in metabolic profiles^58–61^. In a separate cohort of male *TLR4*^flx/flx^, CX3CR1^CreER +/−^ mice that were not subjected to tamoxifen treatment, we observed modest but enduring changes in body weight (data not shown), preventing the use of such mice as an additional control group. Among the studied Ctl and TLR4-MKO mice, we observed tamoxifen-induced reductions in the percentage of body fat in both males and females. However, the Ctl and TLR4-MKO groups otherwise showed no group differences in body weight prior to the experimental period (Figure S2).

In order to investigate the potential role of microglial TLR4 signaling in mediating obesity-induced outcomes, 3 month-old TLR4-MKO and Ctl mice of both sexes began a 16-week exposure to nutrient-matched diets with either 10% (control diet, CD) or 60% (high-fat diet, HFD) two weeks after tamoxifen treatment. In male mice, HFD was associated with similar increases in body weight in TLR4-MKO and Ctl mice (Figure 1A). We found a significant main effect of time (*F* _(2.604, 164.1)_ = 273.3, *p* < 0.001), diet (*F* _(1, 63)_ = 142.6, *p* < 0.001), and an interaction between time and diet (*F* _(2.6, 164)_ = 152.5, *p* < 0.001) on body weight. There was no main effect of genotype or interactions between diet and genotype on measures of body weight. HFD was also associated with increased adiposity (Figure 1B) in male mice. We found a significant main effect of diet (*F* _(1, 64)_ = 382.9, *p* < 0.001) on percent body fat with no main effect of genotype nor interaction between diet and genotype. Similar patterns were found in food intake. There was a significant main effect of diet (*F* _(1, 58)_ = 104.5, *p* < 0.001), such that TLR4-MKO and Ctl male mice fed on HFD had higher daily kilocalorie consumption compared with mice fed on CD (Figure 1C). There was neither a significant main effect of genotype nor an interaction between diet and genotype.

**Figure 1.**
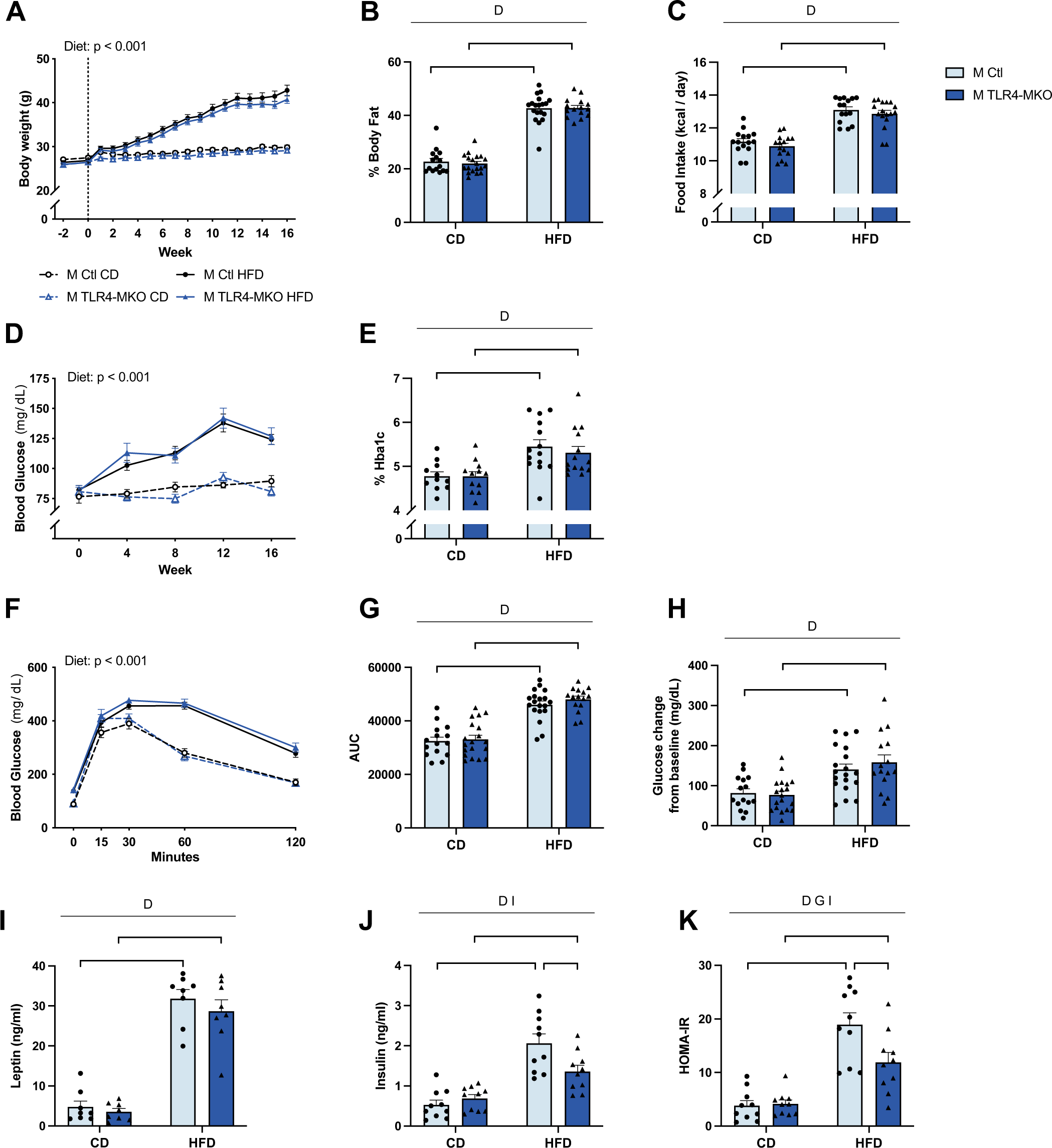
Effects of diet and microglial TLR4 deletion on body weight and metabolic outcomes in male mice. (A) Body weights of control (Ctl) and TLR4-MKO male mice maintained on control diet (CD) or high-fat diet (HFD) during the 16-week dietary treatment; vertical dotted line indicates the week dietary treatment started (n=15-19/group). (B) Percent body fat at the conclusion of the experiment (n=15-19/group). (C) Calculated averages of daily caloric intake (n=15-18/group) across the entire treatment period. (D) Blood glucose levels following 16 h overnight fasting measured at weeks 0, 4, 8, 12, and 16 of the dietary treatment period (n=15-19/group). (E) Levels of blood Hba1c at the end of the dietary treatment (n=11-14/group). (F) Blood glucose levels over time during a 120-minute glucose tolerance test (n=15-19/group). (G) Calculated AUC for the glucose tolerance test (n=15-19/group). (H) Change in fasting blood glucose levels between 0 min (baseline) and 120 min of the glucose tolerance test (n=15-19/group). (I) Plasma leptin (n=8/group) and (J) insulin levels (n=10/group) at the end of the dietary treatment. (K) Calculated insulin resistance (HOMA-IR) scores (n=10/group). Data are presented as mean (+SEM) values that, in some cases, are accompanied by values from individual animals. For figures A, D, and F, values for Ctl mice are shown as black circles and TLR4-MKO mice as blue triangles; control diet are open symbols with dashed line, high-fat diet are filled symbols with solid lines. For panels B-C, E, and G-K, bar graphs are shown in light blue bars for Ctl mice and dark blue bars for TLR4-MKO mice; individual values are shown as filled circles for Ctl mice and filled triangles for TLR4-MKO mice. In panels A, D, and F, statistically significant main effects and associated *p* values are listed. In all other panels, statistically significant main effects are denoted by D (diet), G (genotype) and I (diet and genotype interaction); significant between group comparisons of interest are identified by brackets with * *p* < 0.05, ** *p* < 0.01, *** *p* < 0.001.

To investigate metabolic outcomes of diet-induced obesity, we first measured fasting levels of blood glucose during the 16-week diet period in TLR4-MKO and Ctl male mice. Regardless of genotype, mice fed on HFD showed a significant increase in blood glucose (*F* _(1, 64)_ = 96.0, *p* < 0.001; Figure 1D) at 4-, 8-, 12- and 16-weeks (*p* < 0.05). When comparing glycated hemoglobin (Hba1c) levels in blood, we found a significant main effect of diet (*F* _(1, 64)_ = 382.9, *p* < 0.001; Figure 1E) and no main effect of genotype nor interaction between diet and genotype. Additionally, we performed a glucose tolerance test and mice fed on HFD from both genotypes showed higher levels of blood glucose (*F* _(1, 64)_ = 77.2, *p* < 0.001; Figure 1F) relative to matched CD groups at 60- and 120-min time points (*p* < 0.001). We also calculated AUC (Figure 1G) and change in glucose levels at 120 min from baseline (Figure 1H) to access glucose tolerance. Again, there were significant main effects of diet on AUC (*F* _(1, 64)_ = 102.9, *p* < 001) and change in glucose (*F* _(1, 64)_ = 28.1, *p* < 0.001) whereby HFD mice performed significantly poorer than mice fed on CD on glucose clearance. No significant main effect of genotype or interactions between diet and genotype were observed on AUC and change in glucose measurements.

To further examine metabolic consequences of HFD in the context of microglial TLR4 deletion in male mice, we assessed plasma levels of leptin and insulin following an overnight fast. HFD resulted in significantly increased plasma leptin (*F* _(1, 28)_ = 171.3, *p* < 0.001, Figure 1I) and insulin (*F* _(1, 36)_ = 48.9, *p* < 0.001; Figure 1J) in both TLR4-MKO and Ctl mice. Interestingly, we also found significant interactions between diet and genotype treatment on insulin levels (*F* _(1, 36)_ = 7.5, p = 0.010) in which microglial TLR4 deletion was associated with significant attenuation of the HFD-induced increase in plasma insulin, an effect not observed under CD. We also assessed effects of HFD on insulin resistance by calculating HOMA-IR (Figure 1K). There was a significant main effect of diet (*F* _(1, 36)_ = 55.7, *p* < 0.001), genotype (*F* _(1, 36)_ = 4.8, *p* = 0.035), and interaction between diet and genotype (*F* _(1, 36)_ = 5.8, *p* = 0.021) on HOMA-IR, such that TLR4-MKO mice were less insulin resistant compared with Ctl mice when fed HFD.

**Figure S1.**
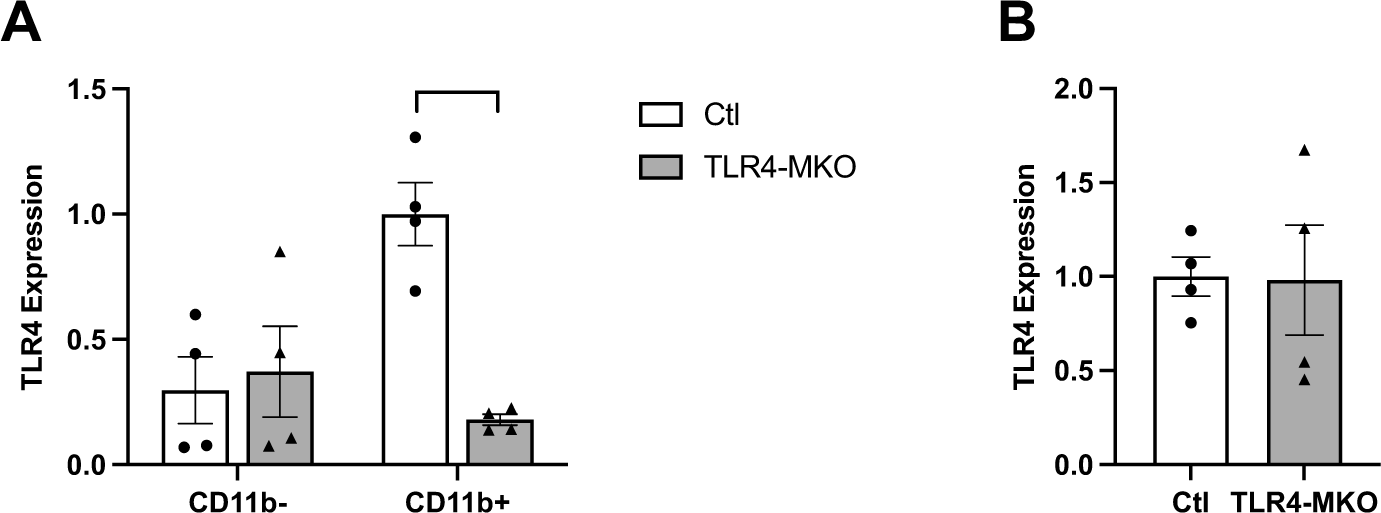
Validation of TLR4 deletion efficacy in TLR4-MKO mice. Quantitative real-time PCR was used to measure mRNA expression levels of TLR4. (A) Brain cells were isolated from control (Ctl, open bars) and TLR4-MKO (gray bars) mice three weeks following tamoxifen injections, then separated by flow cytometry into populations expressing (CD11b+) or not expressing (CD11b-) the microglia/macrophage-specific factor CD11b. (B) Peripheral monocytes collected and analyzed from the same mice. Data show relative TLR4 mRNA expression levels presented as mean (±SEM) values, alongside individual values from each animal; n=4/group, mixed sexes. Brackets identify significant between-group comparisons, with ** *p* < 0.01.

**Figure S2.**
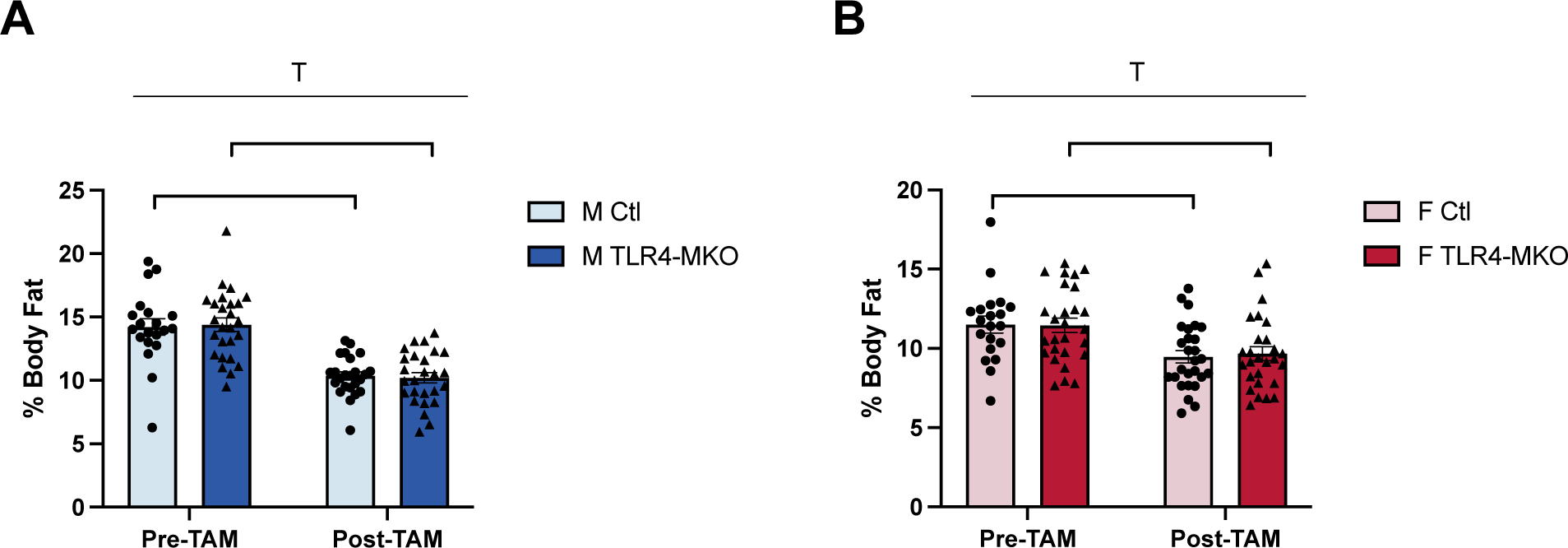
Effects of tamoxifen treatment on adiposity. Body fat percentages were measured prior to (Pre-TAM) and two weeks after (Post-TAM) tamoxifen treatment in control (Ctl) and TLR4-MKO mice in both (A) males (M; Ctl in light blue bars, TLR4-MKO in dark blue) and (B) females (F; Ctl mice in pink bars, TLR4-MKO mice in red bars). Data are presented as mean (±SEM) values and as individual values from each animal (filled circles for Ctl, filled triangles for TLR4-MKO); n=20-27/group. Statistically significant main effects of tamoxifen treatment are denoted by T; significant between group comparisons of interest are identified by brackets with **** *p* < 0.0001.

### 3.2 Microglia TLR4 deletion improved HFD-associated metabolic outcomes in female mice

In females, HFD was also associated with increases in body weight (*F* _(1, 58)_ = 29.8, *p* < 0.001; Figure 2A). Additionally, we found a significant main effect of genotype (*F* _(1, 58)_ = 9.1, *p* = 0.004) and interactions among time, diet and genotype (*F* _(16, 928)_ = 2.6, *p* = 0.01) on body weight during the 16-week treatment period. Between group comparisons revealed that TLR4-MKO mice had significantly lower body weight compared with Ctl mice only when fed HFD (*p* = 0.006). HFD significantly increased percent body fat (*F* _(1, 56)_ = 101.5, *p* < 0.001; Figure 2B) in TLR4-MKO and Ctl mice, and there was a non-significant trend towards an effect of genotype (*p* = 0.053), in which TLR4 deletion was associated with lower percentage body fat in females. The observed differences in body weight and adiposity were likely related to food intake (Figure 2C). There were significant main effects of diet (*F* _(1, 57)_ = 53.9, *p* < 0.001) and genotype (*F* _(1, 57)_ = 6.7, *p* = 0.021) on daily kilocalorie consumption. *Post hoc* tests showed a non-significant trend towards TLR4-MKO mice having lower food intake of HFD compared with Ctl mice (*p* = 0.088).

**Figure 2.**
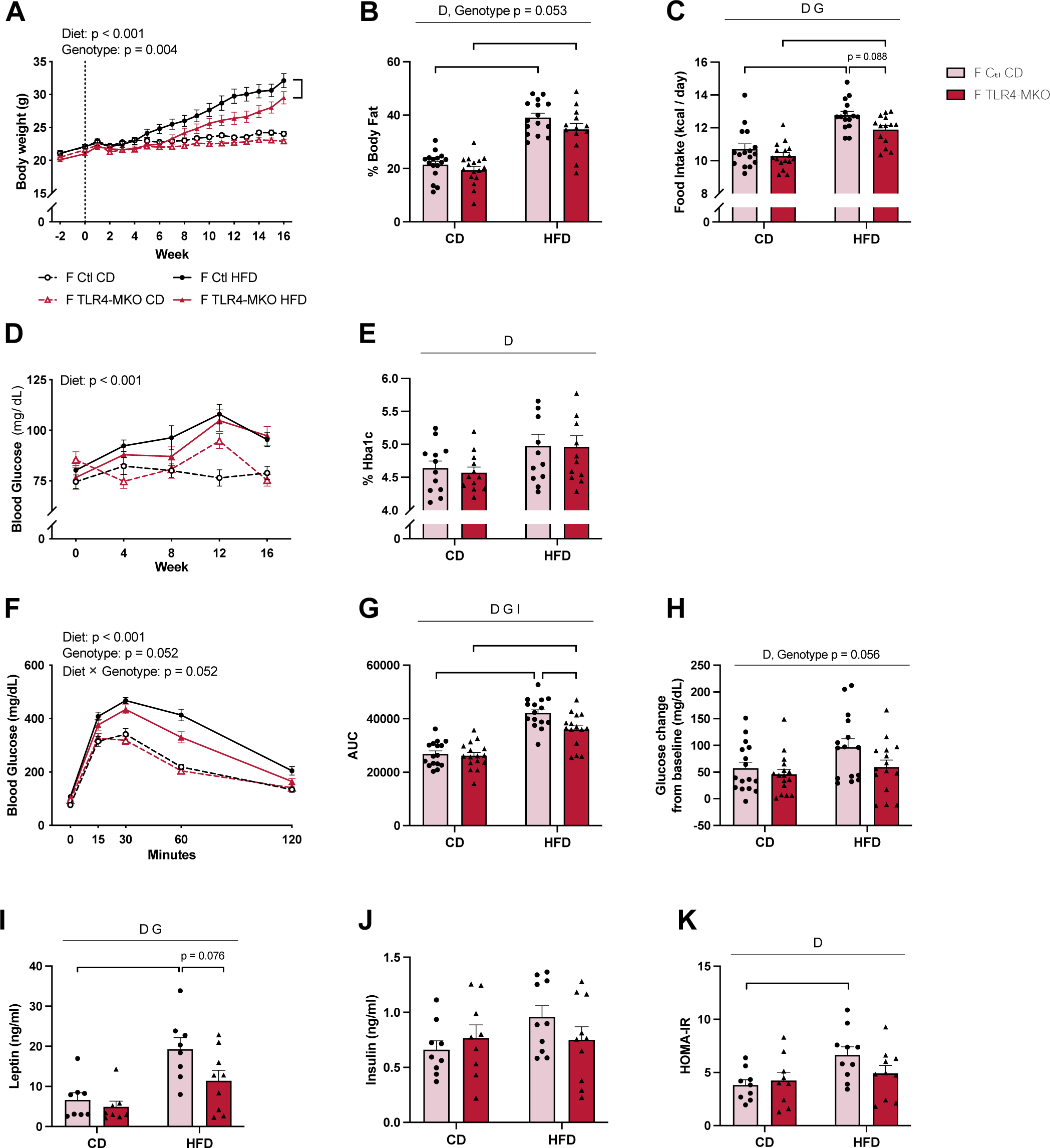
Effects of diet and microglial TLR4 deletion on body weight and metabolic outcomes in female mice. (A) Body weights of control (Ctl) and TLR4-MKO female mice maintained on control diet (CD) or high-fat diet (HFD) during the 16-week dietary treatment; vertical dotted line indicates the week dietary treatment started (n=15-16/group). (B) Percent body fat at the conclusion of the experiment (n=13-16/group). (C) Calculated averages of daily caloric intake (n=14-16/group) across the entire treatment period. (D) Blood glucose levels following 16 h overnight fasting measured at weeks 0, 4, 8, 12, and 16 of the dietary treatment period (n=15-16/group). (E) Levels of blood Hba1c at the end of the dietary treatment (n=12/group). (F) Blood glucose levels over time during a 120-minute glucose tolerance test (n=15-16/group). (G) Calculated AUC for the glucose tolerance test (n=15-16/group). (H) Change in fasting blood glucose levels between 0 min (baseline) and 120 min of the glucose tolerance test (n=15-16/group). (I) Plasma leptin (n=8/group) and (J) insulin levels (n=9-10/group) at the end of the dietary treatment. (K) Calculated insulin resistance (HOMA-IR) scores (n=9-10/group). Data are presented as mean (+SEM) values that, in some cases, are accompanied by values from individual animals. For figures A, D, and F, values for Ctl mice are shown as black circles and TLR4-MKO mice as red triangles; control diet are open symbols with dashed line, high-fat diet are filled symbols with solid lines. For panels B-C, E, and G-K, bar graphs are shown in pink bars for Ctl mice and red bars for TLR4-MKO mice; individual values are shown as filled circles for Ctl mice and filled triangles for TLR4-MKO mice. In panels A, D, and F, statistically significant main effects and associated *p* values are listed. In all other panels, statistically significant main effects are denoted by D (diet), G (genotype) and I (diet and genotype interaction); significant between group comparisons of interest are identified by brackets with * *p* < 0.05, ** *p* < 0.01, *** *p* < 0.001.

The differences among groups in body weight and adiposity were paralleled by the effects of diet and TLR4 deletion on metabolic measures. HFD was associated with a significant increase in fasting glucose levels (*F* _(1, 58)_ = 21.9, *p* < 0.001; Figure 2D) in females. Between group comparisons showed HFD increased fasting glucose measured at weeks 8, 12 and 16 (*p* < 0.05) in Ctl mice, whereas TLR4-MKO mice maintained on HFD showed higher fasting glucose at weeks 4 and 16 (*p* < 0.05) relative to CD. HFD also increased plasma Hba1C levels in mice of both genotypes (*F* _(1, 44)_ = 6.9, *p* = 0.012; Figure 2E). No significant main effect of genotype and no interactions between diet and genotype were found. When challenging the mice to a glucose tolerance test, we found a significant main effect of diet (*F* _(1, 58)_ = 76.04, *p* < 0.001), and non-significant trends towards a main effect of genotype (*p* = 0.052) and an interaction between diet and genotype (*p* = 0.052) on blood glucose (Figure 2F). Between group comparisons revealed that in Ctl mice HFD resulted in higher blood glucose relative to CD at all time points (*p* < 0.001), whereas in TLR4-MKO mice the significant HFD-induced glucose elevation was only observed at the 30 and 60 min time points (*p* < 0.001). Analysis of GTT AUC showed significant main effects of diet (*F* _(1, 58)_ = 82.8, *p* < 0.001) and genotype (*F* _(1, 58)_ = 6.1, *p* = 0.016) as well as an interaction between diet and genotype (*F* _(1, 58)_ = 4.1, *p* = 0.049) such that HFD increased AUC in both genotypes and microglial TLR4 deletion reduced AUC only in the HFD condition (Figure 2G). Additionally, female mice fed HFD were impaired at returning to baseline glucose levels compared with CD-fed mice (*F* _(1, 58)_ = 4.4, *p* = 0.040; Figure 2H) with a non-significant trend towards TLR4-MKO mice performing better than Ctl mice (*p* = 0.056) (Figure 2H).

When examining fasting levels of plasma leptin in females, we found significant main effects of diet (*F* _(1, 29)_ = 18.4, *p* < 0.001) and genotype (*F* _(1, 29)_ = 4.6, *p* = 0.039), though *post hoc* tests showed that the HFD-induced increase was significant only in Ctl mice (*p* = 0.002) (Figure 2I). Neither HFD nor microglial TLR4 deletion significantly affected fasting insulin levels (Figure 2J). However, there was a significant main effect of diet in calculated HOMA-IR (*F* _(1, 34)_ = 6.1, *p* = 0.019; Figure 2K) such that HFD induced insulin resistance only in Ctl mice (*p* = 0.038).

### 3.3 Microglial TLR4 deletion reduced HFD-associated peripheral inflammation

DIO is characterized by low-grade chronic inflammation that affects many organs including adipose tissue^62^. While the expression of CX3CR1 is primarily limited to a small subset of macrophages within the adipose tissue ^42^, it is noteworthy that HFD has been reported to induce an expansion of the adipose macrophage pool^63^. This expansion results from an interplay of factors, including local proliferation^64,65^ as well as the infiltration of peripheral macrophage^66^ and other immune cell types^67^. Therefore, we first assessed TLR4 expression in the visceral fat pad, which allows us to address microglial specificity but also acknowledges the potential influence of peripheral factors on the observed effects. In male mice, neither diet nor genotype significantly affected TLR4 expression levels in adipose tissue (Figure 3A). Next, we examined expressions of inflammatory cytokines at the mRNA level. In males, there was a significant main effect of diet on TNFα (*F* _(1, 28)_ = 4.790, *p* = 0.037; Figure 3B) with *post hoc* tests showing that HFD significantly increased TNFα expression in Ctl (*p* = 0.038) but not TLR4-MKO mice. Further, there were significant interactions between diet and genotype on expressions of IL-1β (*F* _(1, 28)_ = 8.87, *p* = 0.006; Figure 3C) and IL-6 (*F* _(1, 28)_ = 6.24, *p* = 0.019; Figure 3D). Between-group comparisons revealed that HFD increased IL-β and IL-6 levels in Ctl mice (*p* < 0.05), whereas HFD was associated with decrease expression of IL-6 only in TLR4-MKO mice (*p* = 0.001). For the macrophage markers CD68 and F4/80, we observed a significant main effect of diet on CD68 expression (*F* _(1, 28)_ = 36.71, *p* < 0.001; Figure 3E) in which HFD increased CD68 expression levels in both Ctl and TLR4-MKO mice (*p* < 0.01). There was also a main effect of diet on F4/80 levels (*F* _(1, 28)_ = 5.462, *p* = 0.027; Figure 3F), though between group analyses did not reach statistical significance in either Ctl or TLR4-MKO mice. Finally, we examined expression of MHCII protein H2ab1 and MHCII invariant chain peptide CD74. There were significant main effects of diet on levels of H2ab1 (*F* _(1, 28)_ = 6.9, *p* = 0.014; Figure 3G) and CD74 (*F* _(1, 28)_ = 4.73, *p* = 0.038; Figure 3H). Between-group comparisons revealed that HFD is associated with increased H2ab1 and CD74 expression in Ctl mice but not TLR4-MKO mice.

**Figure 3.**
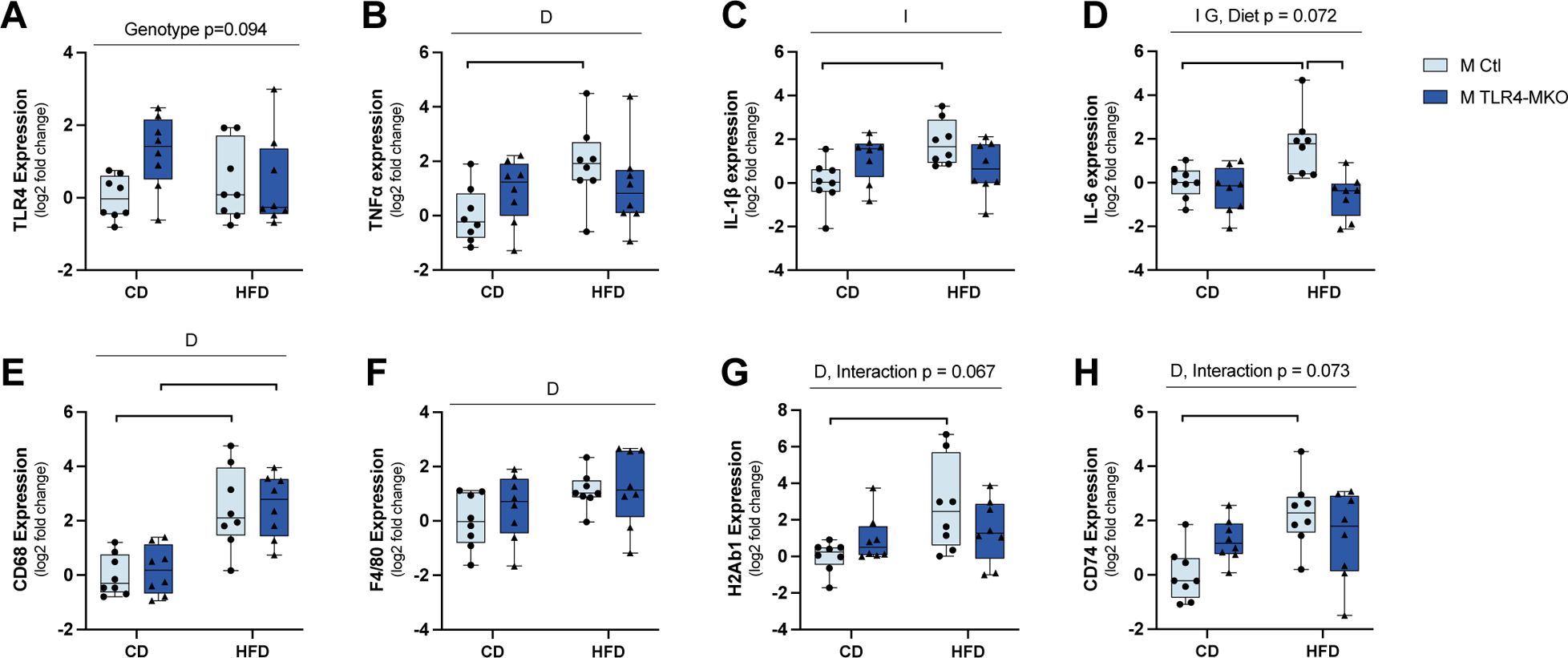
Effects of diet and microglial TLR4 deletion on adipose inflammation in male mice. Quantitative real-time PCR was used to quantify mRNA expression levels in visceral adipose tissue from male control (Ctl, light blue bars) and TLR4-MKO (dark blue bars) mice following exposure to control (CD) or high-fat (HFD) diets. Assessed gene targets were (A) TLR4; cytokines (B) tumor necrosis factor-α (TNFα), (C) interleukin-1β (IL-1β) and (D) interleukin-6 (IL-6); macrophages markers (E) cluster of differentiation 68 (CD68) and (F) EGF-like module-containing mucin-like hormone receptor-like 1 (F4/80); and MHCII-related markers (G) histocompatibility 2, class II antigen A, beta 1 (H2Ab1) and (H) HLA class II histocompatibility antigen gamma chain (CD74). Data are represented as log2 fold change, with quantiles, brackets showing minimum and maximum values, and including individual values (filled circles for Ctl, filled triangles for TLR4-MKO) from each animal; n=8/group. Statistically significant main effects are denoted by D (diet), G (genotype) and I (diet and genotype interaction); significant between group comparisons of interest are identified by brackets with * *p* < 0.05, ** *p* < 0.01, *** *p* < 0.001.

In females, HFD significantly increased TLR4 expression in the adipose tissue (*F* _(1, 28)_ = 4.511, *p* = 0.043; Figure 4A). Among the inflammatory cytokines and macrophages markers, there was a general pattern of HFD-associated increases that was observed for levels of TNFα (*F* _(1, 28)_ = 10.0, *p* = 0.004; Figure 4B), IL-6 (*F* _(1, 28)_ = 7.9, *p* = 0.009; Figure 4D), F4/80 (*F* _(1, 28)_ = 6.1, *p* = 0.020; Figure 4F), H2ab1 (*F* _(1, 28)_ = 6.2, *p* = 0.020; Figure 4G) and CD74 (*F* _(1, 28)_ = 8.0, *p* = 0.009; Figure 4H). For H2ab1, *post hoc* tests showed a statistically non-significant trend towards an HFD-induced increased H2ab1 only in Ctl mice. Microglial TLR4 deletion did not significantly alter mRNA levels of these genes; there was a non-significant trend of lower IL-6 in TLR-MKO mice. We found significant interactions between diet and genotype on levels of CD68 (*F* _(1, 28)_ = 4.8, *p* = 0.038; Figure 4E) and a non-significant trend towards an effect of diet (*p* = 0.055) in which HFD increased CD68 expression levels only on Ctl mice (*p* = 0.030). Diet and genotype did not significantly affect IL-1β expression (Figure 4C).

**Figure 4.**
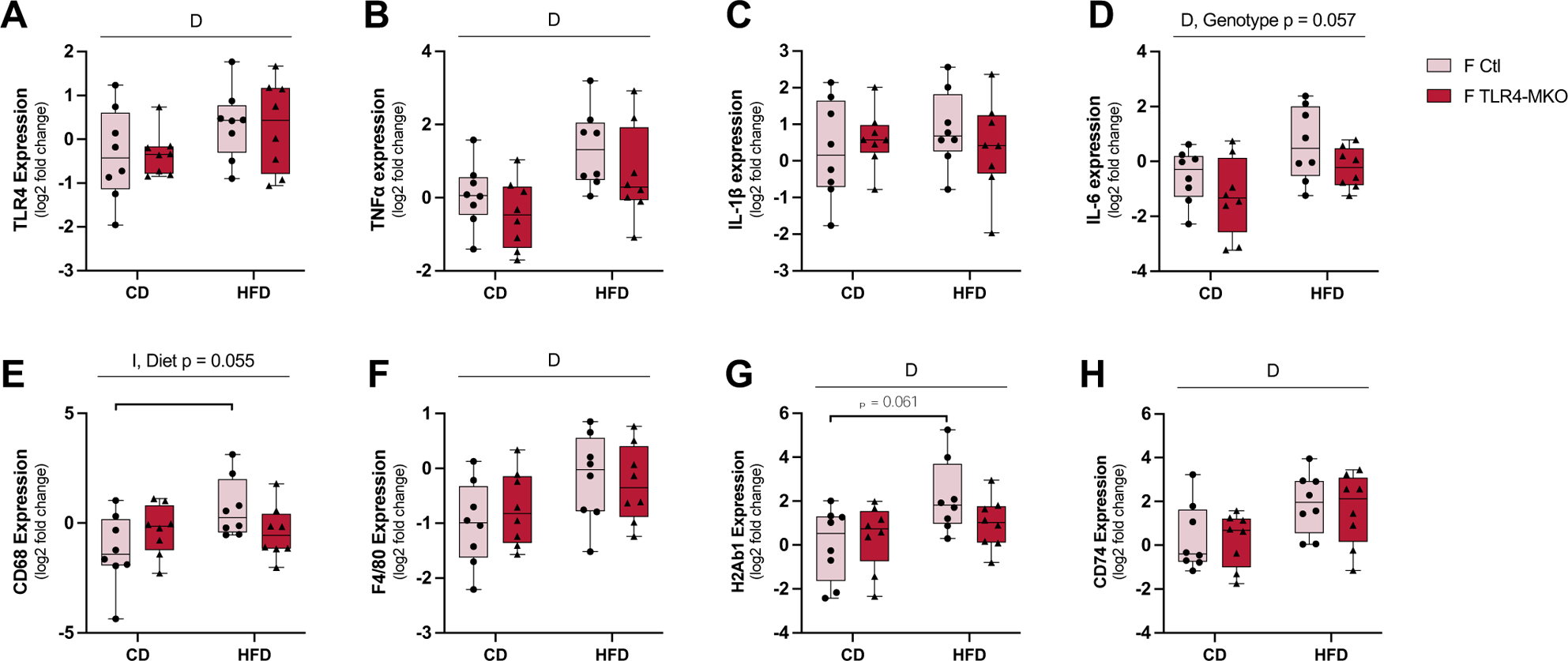
Effects of diet and microglial TLR4 deletion on adipose inflammation in female mice. Quantitative real-time PCR was used to quantify mRNA expression levels in visceral adipose tissue from female control (Ctl, pink bars) and TLR4-MKO (red bars) mice following exposure to control (CD) or high-fat (HFD) diets. Assessed gene targets were (A) TLR4; cytokines (B) TNFα, (C) IL-1β and (D) IL-6; macrophages markers (E) CD68 and (F) F4/80; and MHCII-related markers (G) H2Ab1 and (H) CD74. Data are represented as log2 fold change, with quantiles, brackets showing minimum and maximum values, and including individual values (filled circles for Ctl, filled triangles for TLR4-MKO) from each animal; n=8-9/group. Statistically significant main effects are denoted by D (diet) and I (diet and genotype interaction); significant between group comparisons of interest is identified by brackets with * p < 0.05.

### 3.4 Effects of HFD and microglial TLR4 deletion on plasma cytokine levels

We assessed levels of circulating cytokines in plasma as another measure of peripheral inflammation. In male mice, HFD was associated with a modest effect on plasma levels of cytokines with significant main effects of diet on TNFα (*F* _(1, 36)_ = 4.2, *p* = 0.047; Figure 5A), IL-5 (*F* _(1, 36)_ = 26.4, *p* < 0.001; Figure 5C), and KC/GRO (*F* _(1, 36)_ = 8.1, *p* < 0.008; Figure 5G) and a statistically non-significant trend for IFNγ (Figure 5F). There were also limited effects of microglial/macrophage TLR4 deletion on plasma levels cytokines with a significant main effect of genotype (*F* _(1, 36)_ = 4.9, *p* = 0.034; Figure 5D) only on plasma IL-6 levels, characterized by a statistically non-significant trend of TLR4-MKO mice having lower IL-6 than Ctl mice only when fed HFD.

**Figure 5.**
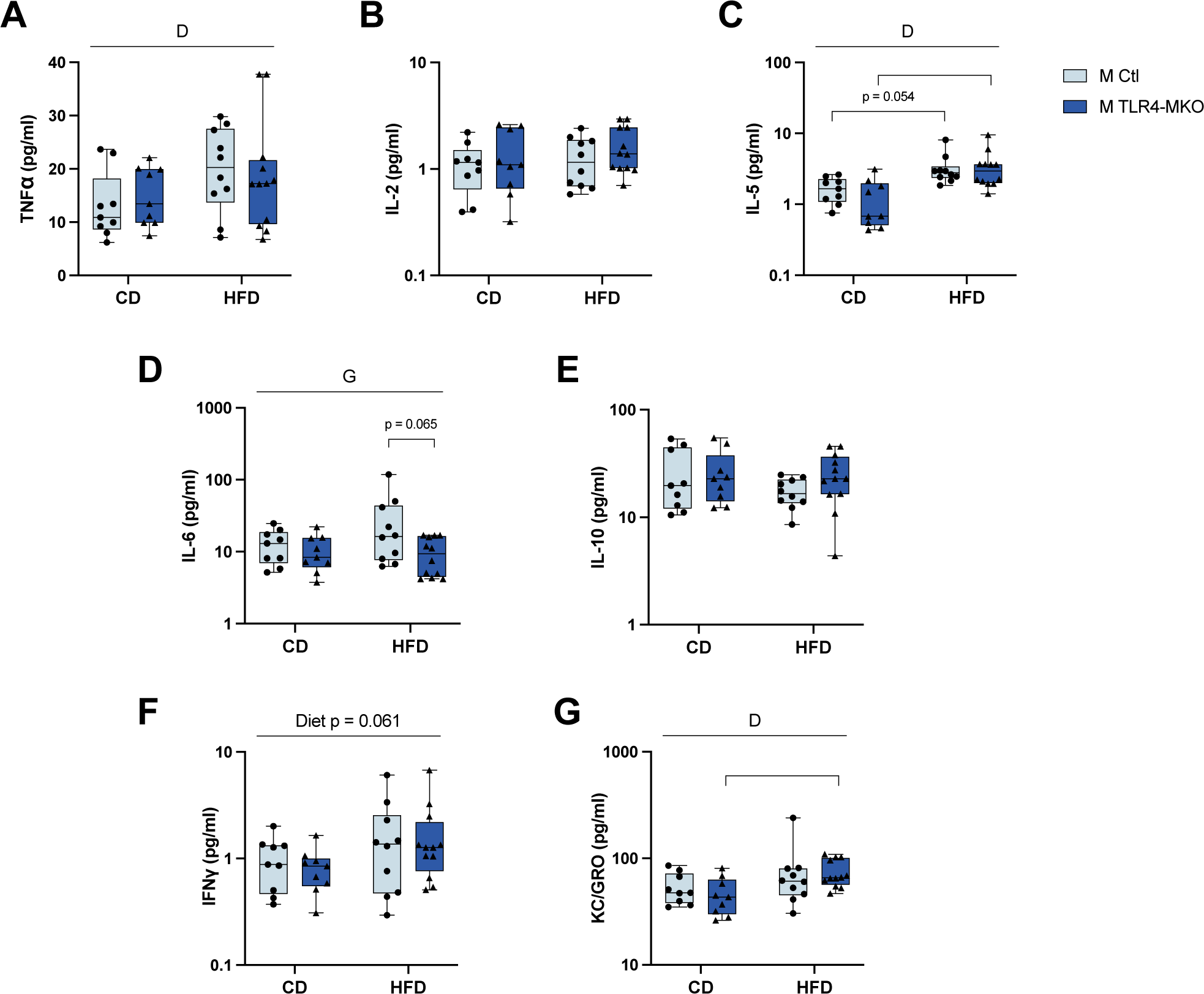
Effects of diet and microglial TLR4 deletion on plasma cytokine levels in male mice. Select inflammation-related factors were measured in plasma from male control (Ctl, light blue bars) and TLR4-MKO (dark blue bars) mice following exposure to control (CD) or high-fat (HFD) diets. Assessed factors were (A) tumor necrosis factor-α (TNFα), (B) interleukin-2 (IL-2), (C) interleukin-5 (IL-5), (D) interleukin-6 (IL-6), (E) interleukin-10 (IL-10), (F) interferon-γ (IFNγ) and (G) keratinocyte chemoattractant/human growth-regulated oncogene (KC/GRO). Data are represented as minimum to maximum values with quantiles, and including individual values (filled circles for Ctl, filled triangles for TLR4-MKO) from each animal; n=9-10/group. Statistically significant main effects are denoted by D (diet) and G (genotype); significant between group comparisons of interest are identified by brackets with * *p* < 0.05, *** *p* < 0.001.

In females, HFD was associated with significantly increased plasma levels of TNFα (*F* _(1, 32)_ = 4.6, *p* = 0.039; Figure 6A), IL-5 (*F* _(1, 32)_ = 28.5, *p* < 0.001; Figure 6C), IL-6 (*F* _(1, 32)_ = 6.2, *p* = 0.018; Figure 6D) and IFNγ (*F* _(1, 32)_ = 5.9, *p* = 0.021; Figure 6F). Additionally, there was a significant main effect of genotype on IL-6 levels (*F* _(1, 32)_ = 6.9, *p* = 0.013) in which microglial/macrophage TLR4 deletion was associated with lower plasma IL-6 levels in the HFD condition (Figure 6D). Genotype was associated with circulating levels of IL-2 (*F* _(1, 32)_ = 4.3, *p* = 0.045; Figure 6B) with a statistically non-significant trend of higher IL-2 in TLR4-MKO HFD-fed mice compared with CD mice.

**Figure 6.**
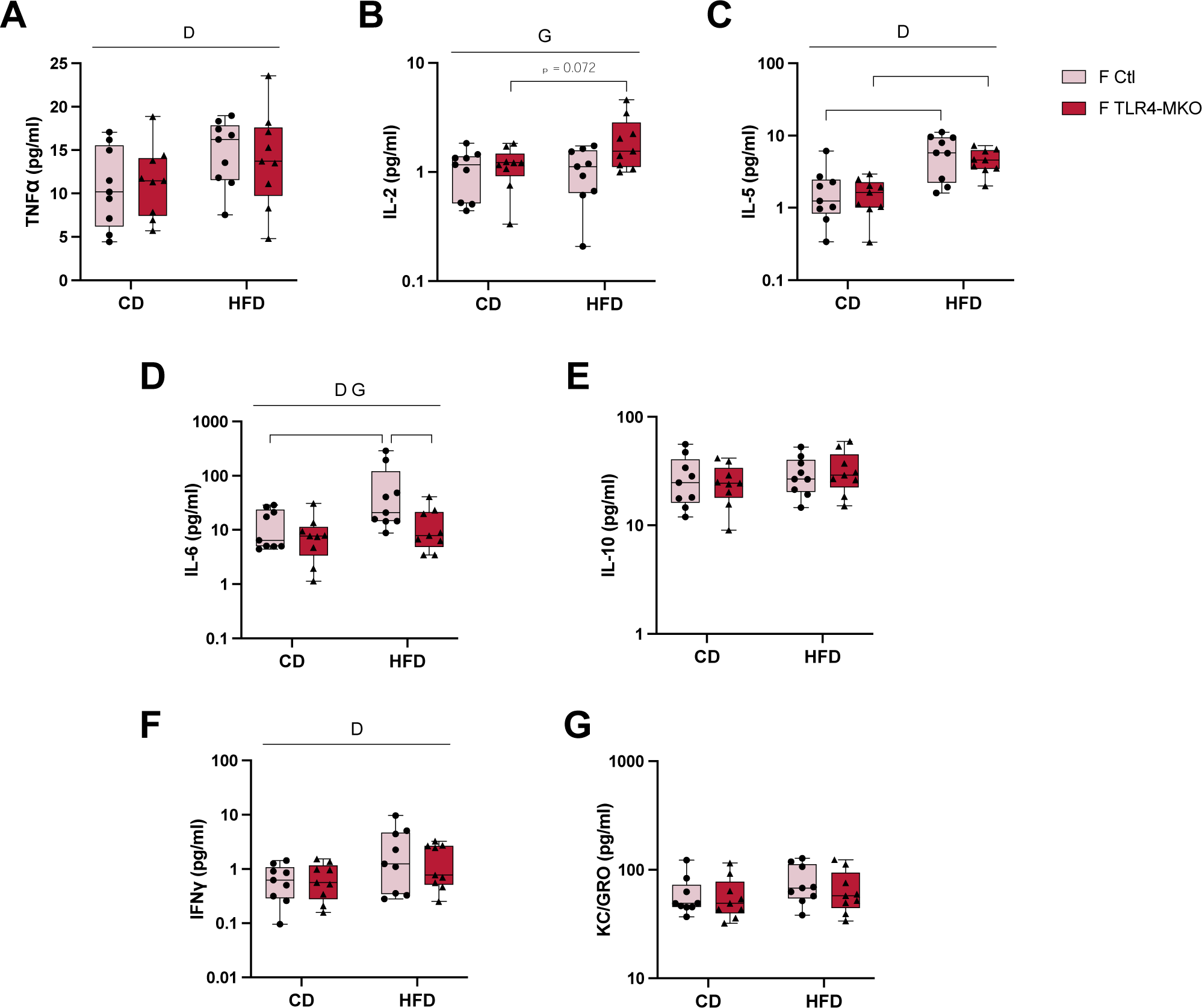
Effects of diet and microglial TLR4 deletion on plasma cytokine levels in female mice. Select inflammation-related factors were measured in plasma from female control (Ctl, pink bars) and TLR4-MKO (red bars) mice following exposure to control (CD) or high-fat (HFD) diets. Assessed factors were (A) tumor necrosis factor-α (TNFα), (B) interleukin-2 (IL-2), (C) interleukin-5 (IL-5), (D) interleukin-6 (IL-6), (E) interleukin-10 (IL-10), (F) interferon-γ (IFNγ) and (G) keratinocyte chemoattractant/human growth-regulated oncogene (KC/GRO). Data are represented as minimum to maximum values with quantiles, and including individual values (filled circles for Ctl, filled triangles for TLR4-MKO) from each animal; n=9 group. Statistically significant main effects are denoted by D (diet) and G (genotype); significant between group comparisons of interest are identified by brackets with * *p* < 0.05, ** *p* < 0.01.

### 3.5 Effects of HFD and microglial TLR4 deletion on gene expression in the hypothalamus

In addition to increasing peripheral inflammation, diet-induced obesity is also associated with a similar type of low-grade inflammation in CNS^36,68^. To investigate the contribution of microglial TLR4 in regulating HFD-induced neuroinflammation, we first assessed gene expression of select cytokines and microglia/macrophage markers in the hypothalamus using quantitative PCR. In male mice, we confirmed that TLR4 expression levels were still significantly lower in TLR4-MKO male mice under both diets after the 16-week experimental period (*F* _(1, 29)_ = 16.2, *p* < 0.001; Figure 7A). For the cytokines, there was a significant main effect of genotype on IL-6 expression (*F* _(1, 29)_ = 6.4, *p* = 0.016; Figure 7B) driven by a statistically non-significant trend of increased IL-6 expression in TLR4-MKO mice on CD. For anti-inflammatory cytokine IL-10, we found a modest but significant interaction between diet and genotype (*F* _(1, 29)_ = 4.9, *p* = 0.035; Figure 7C) in which HFD was associated with increased expression of IL-10 only in Ctl mice. For microglia/macrophage markers, there was a significant main effect of genotype on levels of CD68 (*F* _(1, 29)_ = 4.818, *p* = 0.036; Figure 7D) with TLR4-MKO mice having higher levels of CD68. In contrast, expression of Tmem119, which is specifically expressed by microglia in the brain^69,70^, was increased by HFD only in Ctl mice (Figure 7E). There was a significant main effect of diet (*F* _(1, 29)_ = 6.6, *p* = 0.016) as well as an interaction between diet and genotype (*F* _(1, 29)_ = 7.6, *p* = 0.010) on Tmem119 expression. Trem2 also is expressed in brain largely by microglia^70,71^, however its expression was not significantly different across groups (Figure 7F). Upregulation of MHCII is a marker of reactive microglia^72^, thus we examined gene expression of H2Ab1 and CD74. We found significant main effects of diet on levels of both H2Ab1 (*F* _(1, 29)_ = 4.3, *p* = 0.047; Figure 7G) and CD74 (*F* _(1, 29)_ = 8.4, *p* = 0.008; Figure 7H) with *post hoc* tests revealing non-significant trends of HFD-induced increases in H2Ab1 (*p* = 0.071) and CD74 (*p* = 0.058) specifically in Ctl mice.

**Figure 7.**
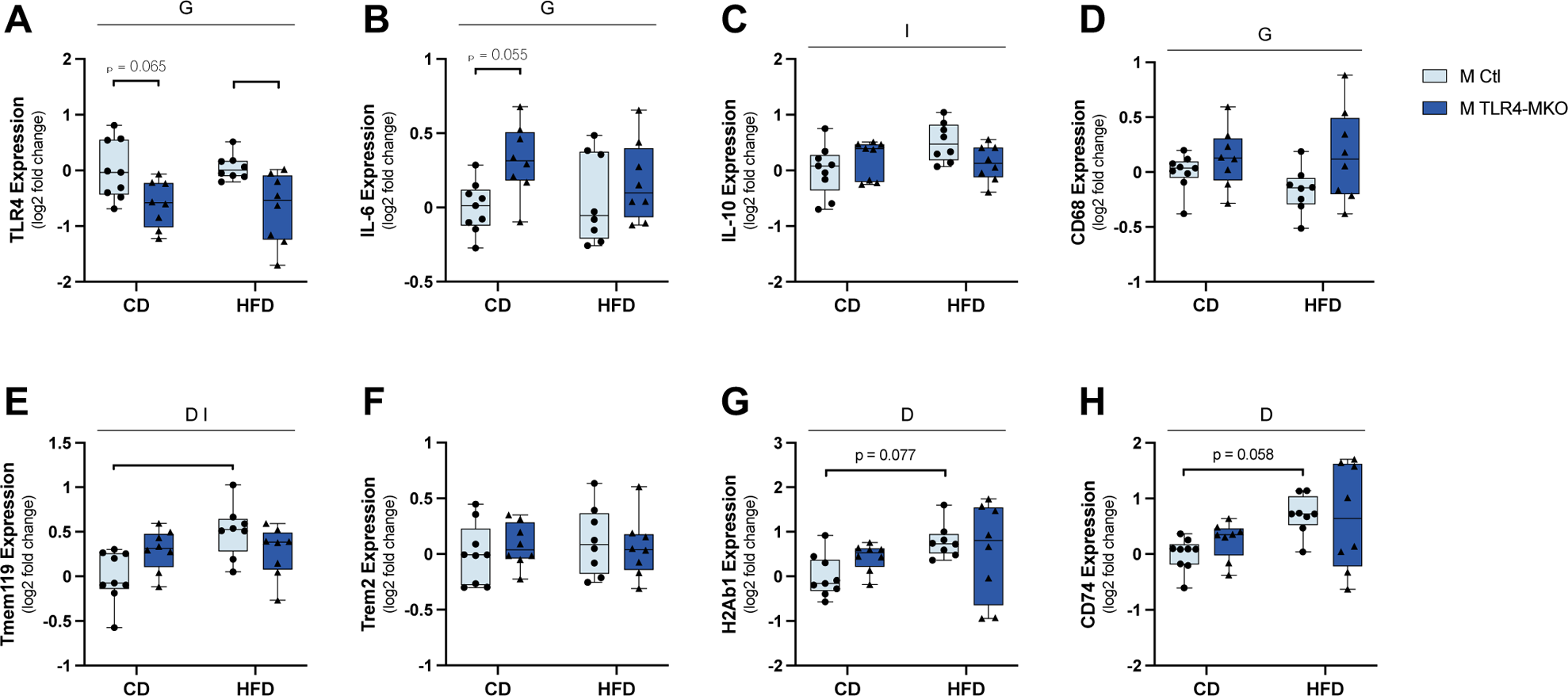
Effects of diet and microglial TLR4 deletion on gene expression in the hypothalamus in male mice. Quantitative real-time PCR was used to quantify mRNA expression levels in hypothalamic brain tissue from male control (Ctl, light blue bars) and TLR4-MKO (dark blue bars) mice following exposure to control (CD) or high-fat (HFD) diets. Assessed gene targets were (A) TLR4; cytokines (B) interleukin-6 (IL-6) and (C) interleukin-10 (IL-10); macrophage markers (D) cluster of differentiation 68 (CD68); microglia marker (E) transmembrane protein 119 (Tmem119) and (F) triggering receptor expressed on myeloid cells 2 (Trem2); and MHCII-related markers (G) histocompatibility 2, class II antigen A, beta 1 (H2Ab1) and (H) HLA class II histocompatibility antigen gamma chain (CD74). Data are represented as log2 fold change, with quantiles, brackets showing minimum and maximum values, and include individual values (filled circles for Ctl, filled triangles for TLR4-MKO) from each animal; n=8-9/group. Statistically significant main effects are denoted by D (diet), G (genotype) and I (diet and genotype interaction); significant between group comparisons of interest are identified by brackets with * *p* < 0.05, ** *p* < 0.01.

Hypothalamic expression of microglia/macrophage markers and cytokines in female mice also showed modest regulation by diet and genotype. As with the males, hypothalamic expression of TLR4 in female mice was significantly affected by genotype (*F* _(1, 29)_ = 7.5, *p* = 0.010; Figure 8A) with TLR4-MKO mice having lower levels than Ctl mice. Moreover, TLR4-MKO mice showed significantly increased IL-6 expression (*F* _(1, 29)_ = 5.5, *p* = 0.026; Figure 8B). We found no significant main effects of diet and genotype and no interactions on the levels of IL-10, CD68, Tmem119 and Trem2 (Figure 8C-F). However, HFD was associated with upregulation of H2Ab1 (*F* _(1, 29)_ = 7.9, *p* = 0.009; Figure 8G) and CD74 (*F* _(1, 29)_ = 5.3, *p* = 0.029; Figure 8H) in the hypothalamus in female mice.

**Figure 8.**
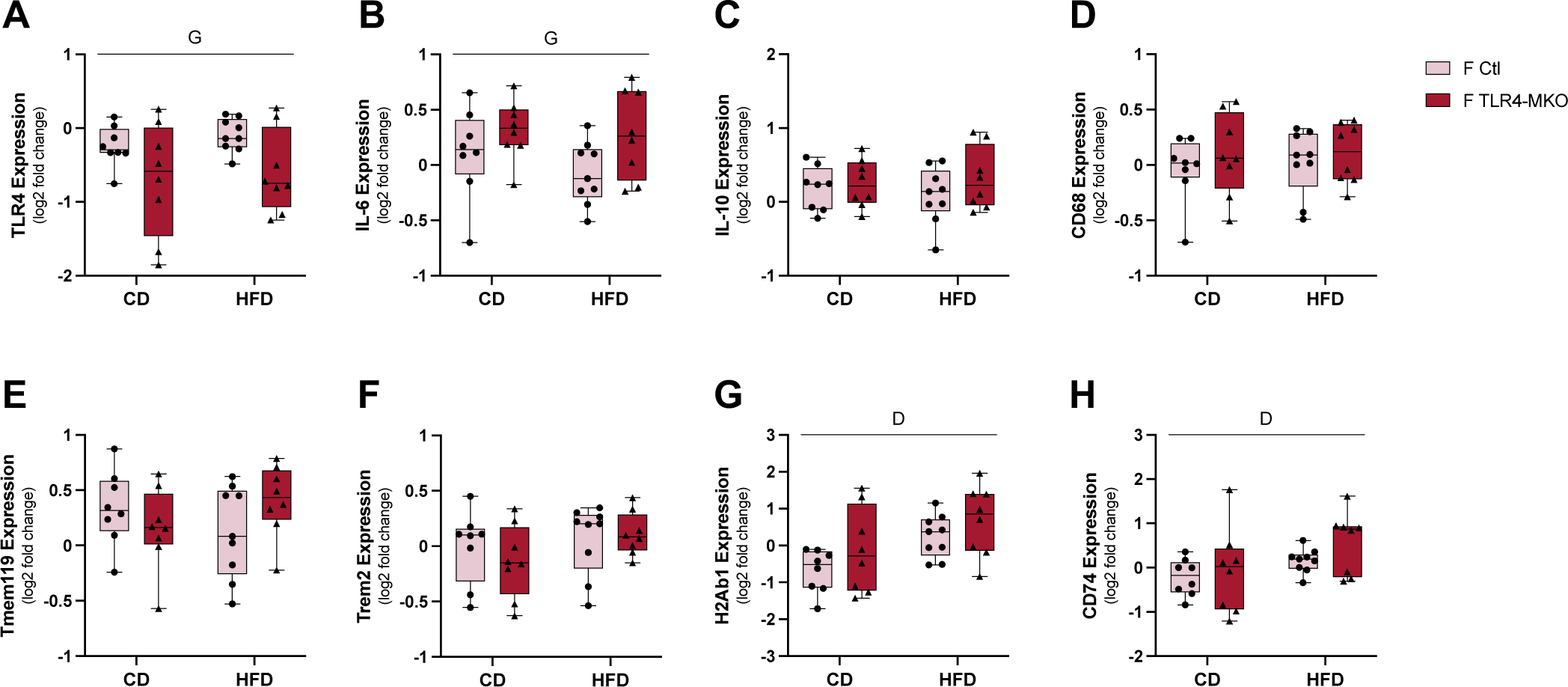
Effects of diet and microglial TLR4 deletion on gene expression in the hypothalamus in female mice. Quantitative real-time PCR was used to quantify mRNA expression levels in hypothalamic brain tissue from male control (Ctl, light blue bars) and TLR4-MKO (dark blue bars) mice following exposure to control (CD) or high-fat (HFD) diets. Assessed gene targets were (A) TLR4; cytokines (B) interleukin-6 (IL-6) and (C) interleukin-10 (IL-10); macrophage markers (D) cluster of differentiation 68 (CD68); microglia marker (E) transmembrane protein 119 (Tmem119) and (F) triggering receptor expressed on myeloid cells 2 (Trem2); and MHCII-related markers (G) histocompatibility 2, class II antigen A, beta 1 (H2Ab1) and (H) HLA class II histocompatibility antigen gamma chain (CD74). Data are represented as log2 fold change, with quantiles, brackets showing minimum and maximum values, and include individual values (filled circles for Ctl, filled triangles for TLR4-MKO) from each animal; n=8-9/group. Statistically significant main effects are denoted by D (diet), and G (genotype).

### 3.6 TLR4-MKO mice were protected from glial activation in the hypothalamus

HFD has been shown to increase activated states of microglia and astrocytes, particularly in the hypothalamus^73^. To address possible contributions of microglial TLR4 signaling to HFD-induced glial activation, we examined activated glial phenotypes in the arcuate nucleus (ARC) in the hypothalamus by (i) analyzing Iba-1+ cell morphology for microglia, and (ii) quantifying GFAP immunoreactivity for astrocytes. In male mice, neither diet nor genotype significantly affected averaged process length and numbers of endpoints in microglia (Figure 9B, C). However, there was a non-significant trend towards an effect of genotype on process length, with TLR4-MKO mice having shorter processes than Ctl mice. Activation phenotypes of microglia are also associated with increased soma size and reduced soma roundness^74^. Upon measuring these complementary measures of microglia activation in male mice, we found a significant main effect of diet (*F* _(1, 20)_ = 5.21, *p* = 0.034) and an interaction between diet and genotype (*F* _(1, 20)_ = 13.41, *p* = 0.002) on microglia soma size (Figure 9D). Between-group comparisons revealed that HFD was associated with larger soma in Ctl mice (*p* = 0.002) that was significantly by microglial TLR4 deletion (*p* = 0.003). There was a non-significant trend of reduced soma roundness by diet (Figure 9E).

**Figure 9.**
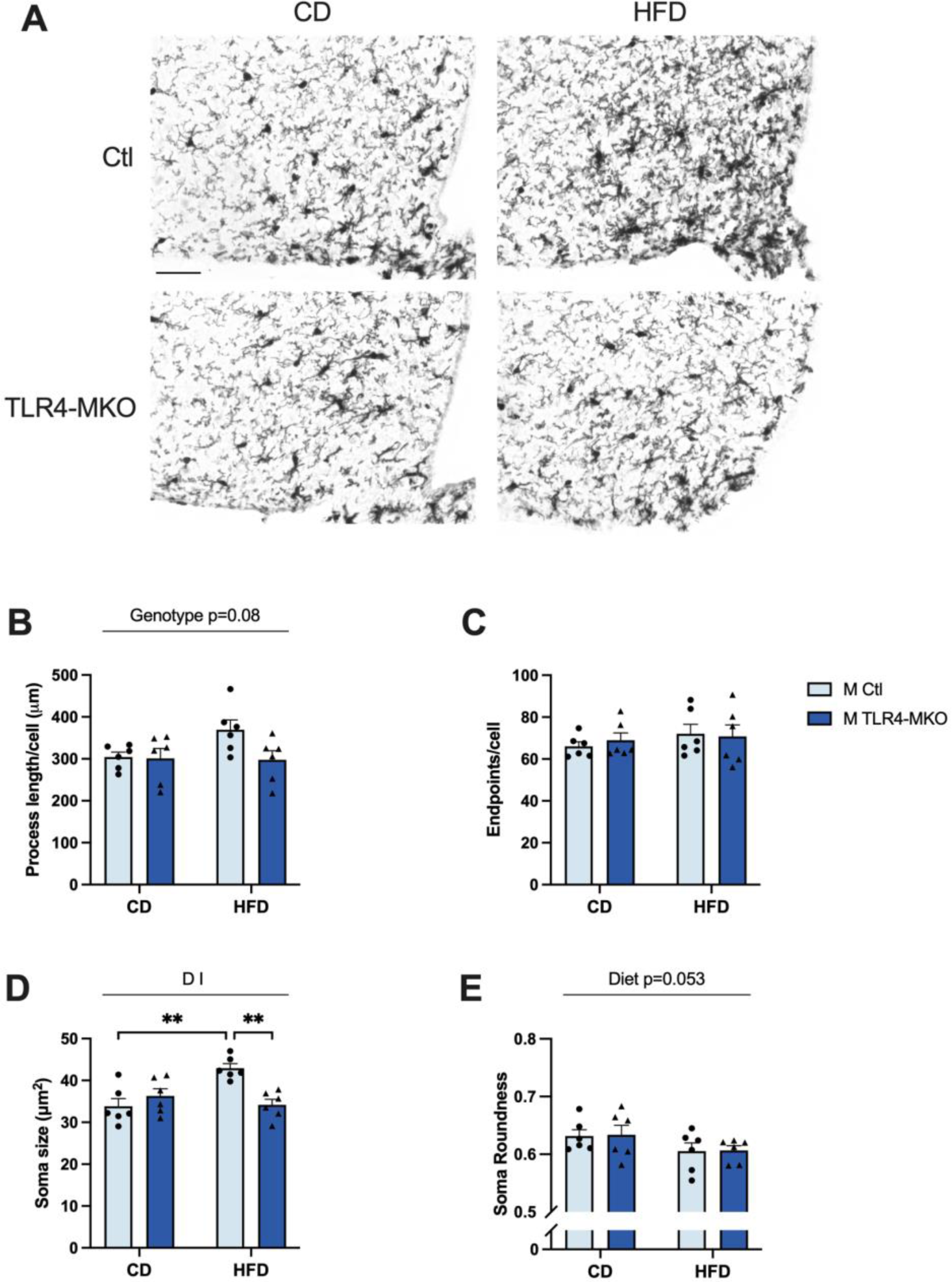
Effects of diet and microglial TLR4 deletion on microglial cell morphology in the hypothalamus in male mice. (A) Representative images of Iba1 immunoreactivity in the arcuate nucleus of hypothalamus from all four male experimental groups. Scale bar = 50 μm. Images of Iba1 immunoreactivity from male control (Ctl, light blue bars) and TLR4-MKO (dark blue bars) mice following exposure to control (CD) or high-fat (HFD) diets were assessed for (B) average process length of Iba1+ cells, (C) average process endpoints per cell of Iba1+ cells, (D) soma size and (E) soma roundness of Iba1+ cells. Data are presented as mean (+SEM) values and include individual values (filled circles for Ctl, filled triangles for TLR4-MKO); n = 6/group. Statistically significant main effects are denoted by D (diet) and I (diet and genotype interaction); significant between group comparisons of interest are identified with brackets and significance levels are denoted by ** *p* < 0.01.

When examining microglial activation across groups in female mice, we found a significant main effect of diet (*F* _(1, 20)_ = 4.9, *p* = 0.039) and an interaction (*F* (_(1, 20)_ = 4.8, *p* = 0.040) on microglia process length in which HFD significantly increased process length only in Ctl mice (Figure 10B). Neither diet nor genotype significantly affected average microglia endpoints (Figure 10C). Microglial soma size showed a significant main effect of genotype (*F* _(1, 20)_ = 4.8, *p* = 0.041) and an interaction between diet and genotype (*F* _(1, 20)_ = 8.2. *p* = 0.009) such that HFD was associated with increased soma size in Ctl but not TLR4-MKO mice (Figure 10D). Lastly, there was a significant main effect of diet on soma roundness (*F* _(1, 20)_ = 5.820, *p* = 0.026; Figure 10E) with Ctl HFD-fed mice having lower roundness compared with mice fed on CD.

**Figure 10.**
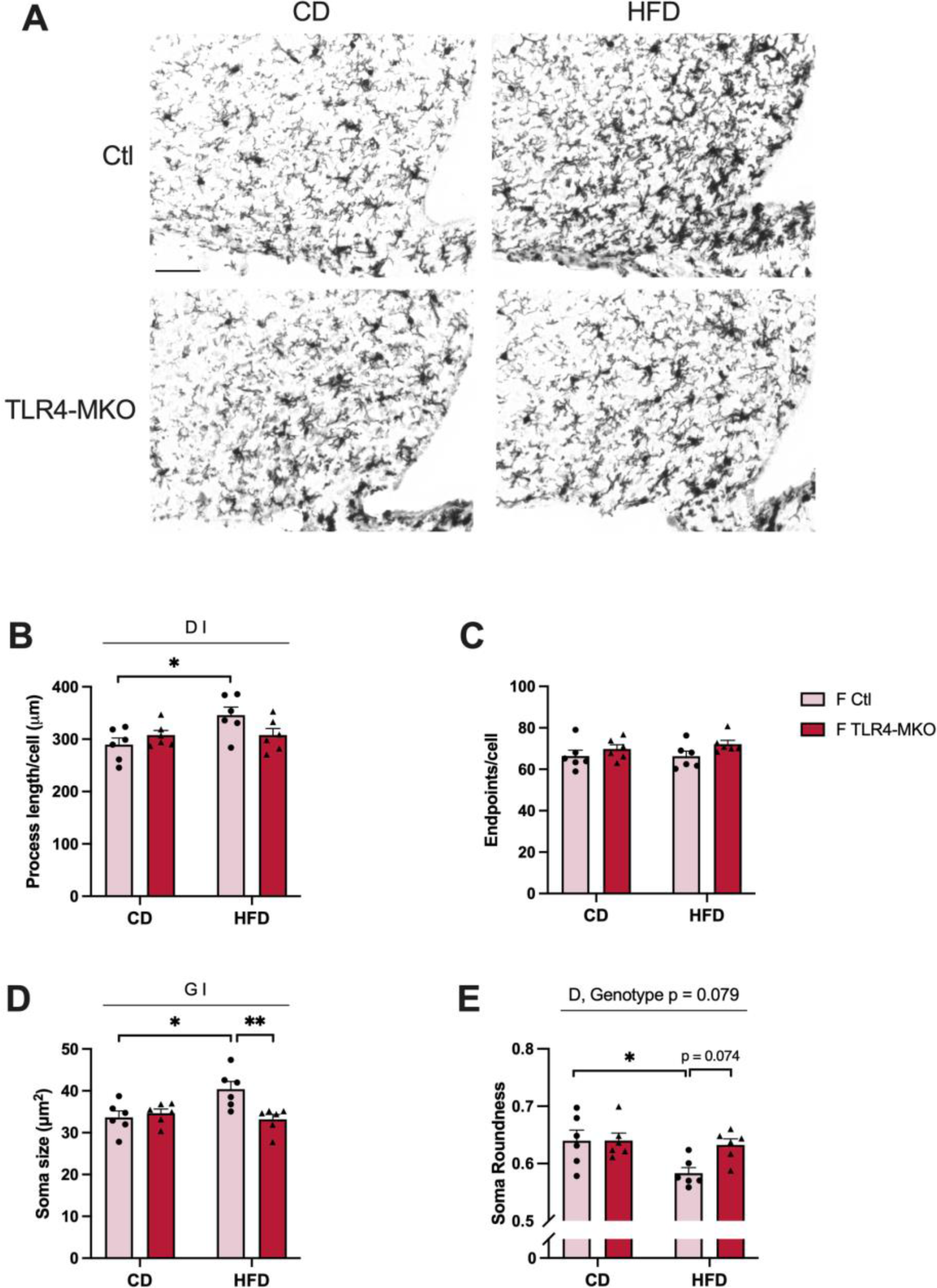
Effects of diet and microglial TLR4 deletion on microglia cell morphology in the hypothalamus in female mice. (A) Representative images of Iba1 immunoreactivity in the arcuate nucleus of the hypothalamus from all four female experimental groups. Scale bar = 50 μm. Images of Iba1 immunoreactivity from female control (Ctl, pink bars) and TLR4-MKO (red bars) mice following exposure to control (CD) or high-fat (HFD) diets were assessed for (B) average process length of Iba1+ cells, (C) average process endpoints per cell of Iba1+ cells, (D) soma size and (E) soma roundness of Iba1+ cells. Data are presented as mean (+SEM) values and include individual values (filled circles for Ctl, filled triangles for TLR4-MKO); n = 6/group. Statistically significant main effects are denoted by D (diet), G (genotype), and I (diet and genotype interaction); significant between group comparisons of interest are identified with brackets and significance levels are denoted by * *p* < 0.05, ** *p* < 0.01.

We next analyzed astrocytes in the ARC of male and female mice. In male mice, the level of GFAP immunoreactivity was significantly increased by HFD (*F* _(1, 20)_ = 7.7, *p* = 0.012), whereas microglial TLR4 deletion was associated with lower levels of astrocyte activation compared with Ctl mice (*F* _(1, 20)_ = 5.1, *p* = 0.036; Figure 11A). In females, HFD induced GFAP immunoreactivity (*F* _(1, 20)_ = 11.0, *p* = 0.004; Figure 11B) but this effect was only significant in Ctl mice.

**Figure 11.**
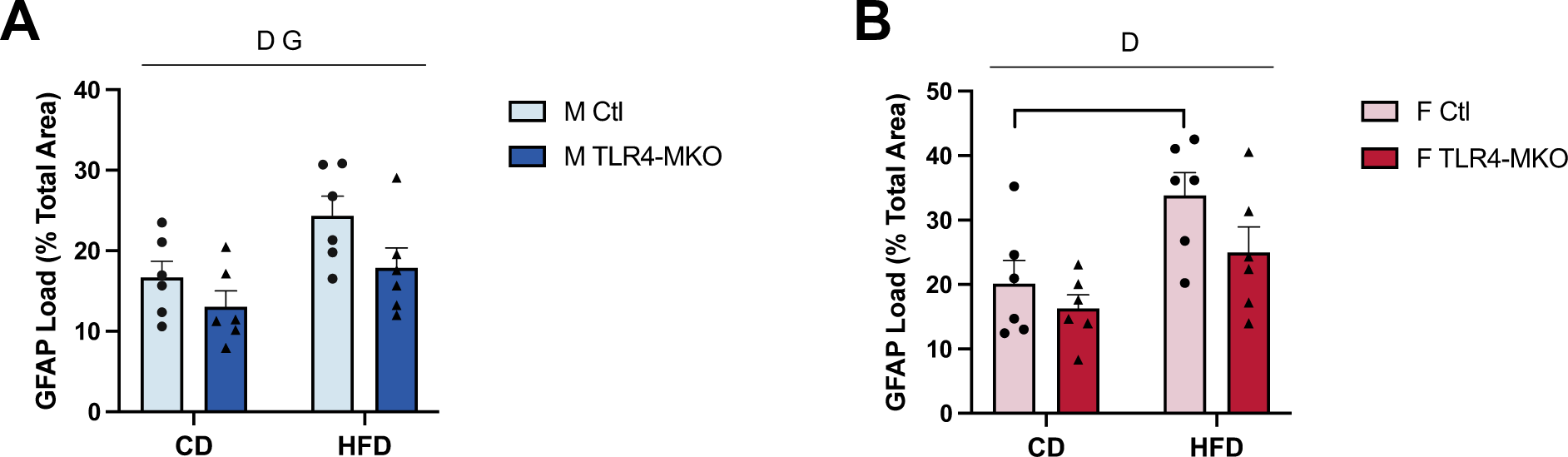
Effects of diet and microglial TLR4 deletion on astrocyte immunoreactivity in the hypothalamus in both sexes. Immunoreactive loads of the astrocyte marker glial fibrillary acidic protein (GFAP) were quantified in the arcuate nucleus of the hypothalamus following exposure to control (CD) or high-fat (HFD) diets in (A) male (Ctl, light blue bars; TLR4-MKO, dark blue bars) and (B) female (Ctl, pink bars; TLR4-MKO, red bars) mice. Data are presented as mean (+SEM) values and include individual values (filled circles for Ctl, filled triangles for TLR4-MKO); n = 6/group. Statistically significant main effects are denoted by D (Diet) and G (Genotype) and significance by * *p* < 0.05.

Together, our data suggested that deletion of macrophage/microglial TLR4 significantly attenuated markers of microglia and astrocyte activation in the hypothalamus in both sexes that results from HFD.

### 3.7 Effects of HFD and microglial TLR4 deletion on hippocampal gene expressions

Neuroinflammation derived from obesity can extend beyond the hypothalamus. Therefore, we assessed gene expression on the same set of markers in the hippocampus by quantitative PCR. In male mice, we observed that TLR4-MKO mice had significant lower TLR4 expression than Ctl mice across both diets (*F* _(1, 31)_ = 32.3, *p* < 0.001; Figure 12A), consistent with a persistent effect of macrophage/microglial TLR4 deletion. Next, results demonstrated a significant interaction between diet and genotype on levels of pro-inflammatory cytokine IL-6 (*F* _(1, 30)_ = 4.8, *p* = 0.037; Figure 12B) with between-group comparisons revealing a non-significant trend toward higher IL-6 levels in TLR4-MKO mice compared with the Ctl mice under CD conditions. However, mRNA levels of anti-inflammatory cytokine IL-10 were significantly upregulated by HFD (*F* _(1, 30)_ = 4.6, *p* = 0.041; Figure 12C) but not affected by genotype. Moreover, no significant diet or genotype effects were observed on expression of the macrophage marker CD68 (Figure 12D). Notably, HFD induced the expression of microglia-specific markers Tmem119 (*F* _(1, 31)_ = 7.4, *p* = 0.011; Figure 12E) and Trem2 (*F* _(1, 31)_ = 4.9, *p* = 0.034; Figure 12F) but the effect reached statistical significance only in Ctl mice. Lastly, genes associated with microglia activation including H2Ab1 (*F* _(1, 30)_ = 5.4, *p* = 0.027; Figure 12G) and CD74 (*F* _(1, 30)_ = 6.7, *p* = 0.015; Figure 12H) were upregulated by HFD, though again the effect was only significant in Ctl mice. Among several markers of neuroinflammation and glial activation, TLR4-MKO mice generally showed attenuated responses to HFD in comparison to Ctl mice.

**Figure 12.**
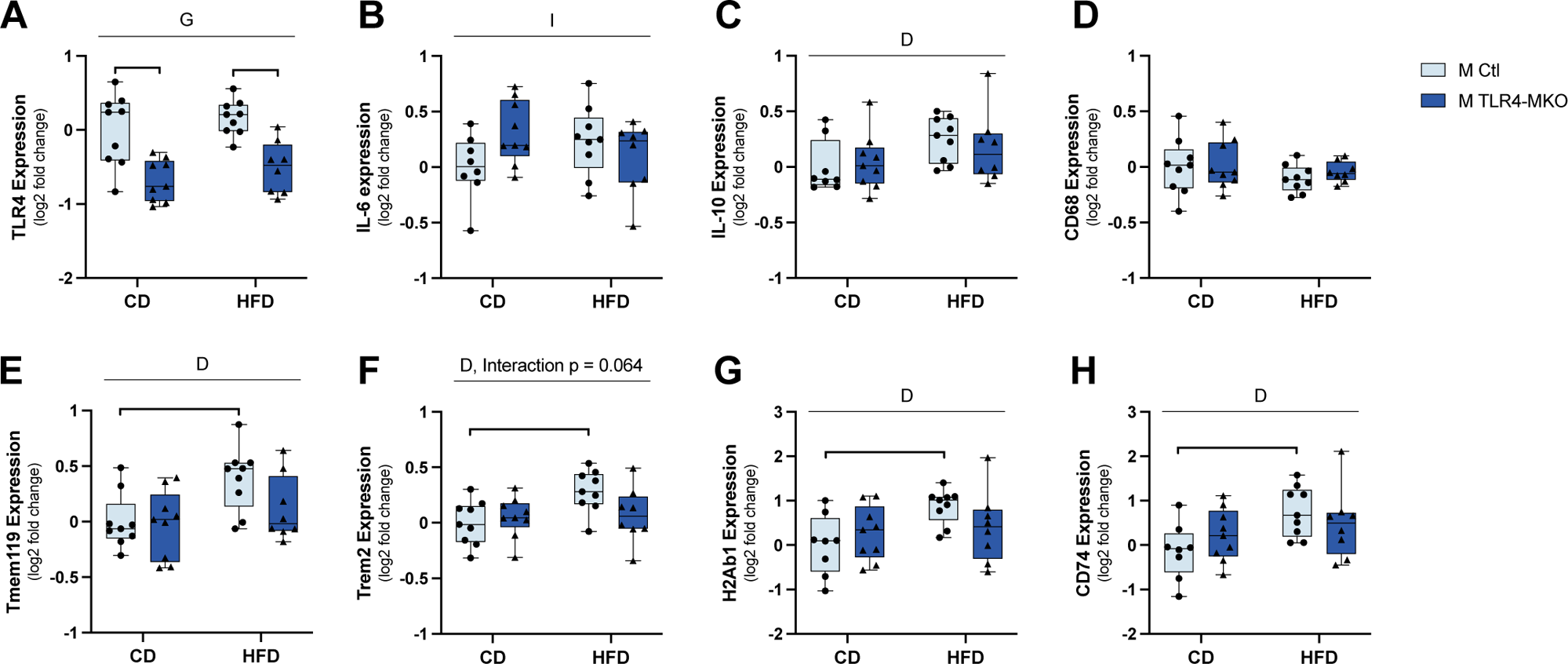
Effects of diet and microglial TLR4 deletion on gene expression in the hippocampus in male mice. Quantitative real-time PCR was used to quantify mRNA expression levels in hippocampal brain tissue from male control (Ctl, light blue bars) and TLR4-MKO (dark blue bars) mice following exposure to control (CD) or high-fat (HFD) diets. Assessed gene targets were (A) TLR4; cytokines (B) interleukin-6 (IL-6) and (C) interleukin-10 (IL-10); macrophage markers (D) cluster of differentiation 68 (CD68); microglia marker (E) transmembrane protein 119 (Tmem119) and (F) triggering receptor expressed on myeloid cells 2 (Trem2); and MHCII-related markers (G) histocompatibility 2, class II antigen A, beta 1 (H2Ab1) and (H) HLA class II histocompatibility antigen gamma chain (CD74). Data are represented as log2 fold change, with quantiles, brackets showing minimum and maximum values, and include individual values (filled circles for Ctl, filled triangles for TLR4-MKO) from each animal; n=8-9/group. Statistically significant main effects are denoted by D (diet), G (genotype) and I (diet and genotype interaction); significant between group comparisons of interest are identified by brackets with * *p* < 0.05, ** *p* < 0.01.

Similar to the males, female TLR4-MKO mice exhibited lower expression levels of TLR4 (*F* _(1, 30)_ = 12.0, *p* = 0.002; Figure 13A) in the hippocampus than Ctl mice. However, we found no significant effects of diet, genotype, or their interaction on expression of any of the other probed genes (Figure 13B-H).

**Figure 13.**
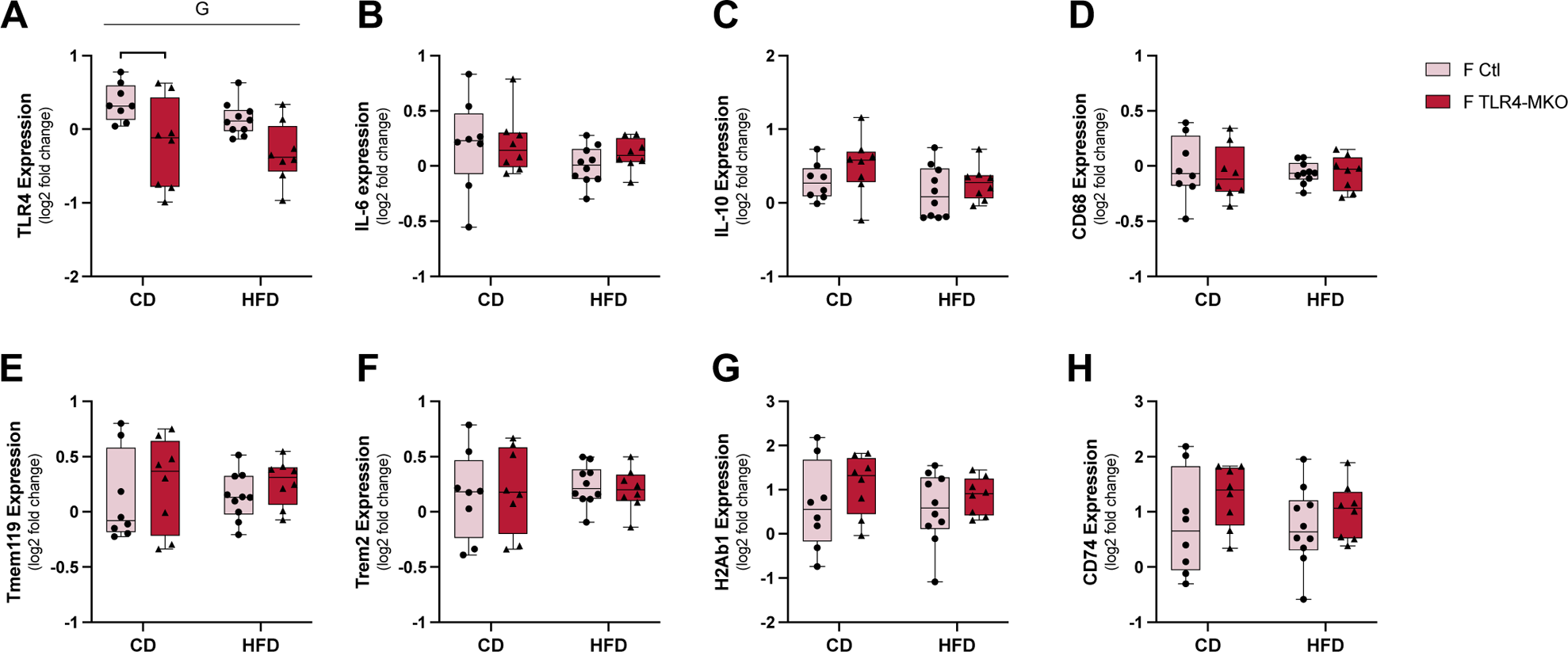
Effects of diet and microglial TLR4 deletion on gene expression in the hippocampus in female mice. Quantitative real-time PCR was used to quantify mRNA expression levels in hypothalamic brain tissue from male control (Ctl, pink bars) and TLR4-MKO (red bars) mice following exposure to control (CD) or high-fat (HFD) diets. Assessed gene targets were (A) TLR4; cytokines (B) interleukin-6 (IL-6) and (C) interleukin-10 (IL-10); macrophage markers (D) cluster of differentiation 68 (CD68); microglia marker (E) transmembrane protein 119 (Tmem119) and (F) triggering receptor expressed on myeloid cells 2 (Trem2); and MHCII-related markers (G) histocompatibility 2, class II antigen A, beta 1 (H2Ab1) and (H) HLA class II histocompatibility antigen gamma chain (CD74). Data are represented as log2 fold change, with quantiles, brackets showing minimum and maximum values, and include individual values (filled circles for Ctl, filled triangles for TLR4-MKO) from each animal; n=8-9/group. Statistically significant main effects are denoted by G (genotype); significant between group comparisons of interest are identified by brackets with * *p* < 0.05.

### 3.8 Male TLR4-MKO mice were protected from glial activation in the hippocampus

As performed in the hypothalamus, we also examined in hippocampus immunohistochemical indices of activated glial phenotypes. In male mice, Iba-1-immunoreactive cells showed a significant main effect of diet on average process length (*F* _(1, 20)_ = 14.2, *p* = 0.001; Figure 14A) and endpoint number (*F* _(1, 20)_ = 5.3, *p* = 0.032; Figure 14B) in which HFD was associated with increased markers of microglial activation only in Ctl mice. Neither diet nor genotype affected microglial soma size in males (Figure 14C). Next, we found a significant interaction between diet and genotype on soma roundness in Iba-1-labeled cells (*F* _(1, 20)_ = 6.5, p = 0.019; Figure 14D) such that TLR4-MKO mice had significantly higher soma roundness than Ctl mice under HFD treatment. Assessment of astrocyte activation in male mice by levels of GFAP immunoreactivity revealed a significant main effect of genotype (*F* _(1, 20)_ = 4.83, p = 0.040; Figure 14E) with lower GFAP immunoreactivity in TLR4-MKO mice. Additionally, there was a non-significant trend towards an effect of diet (*p* = 0.051) moving towards increased GFAP with HFD.

**Figure 14.**
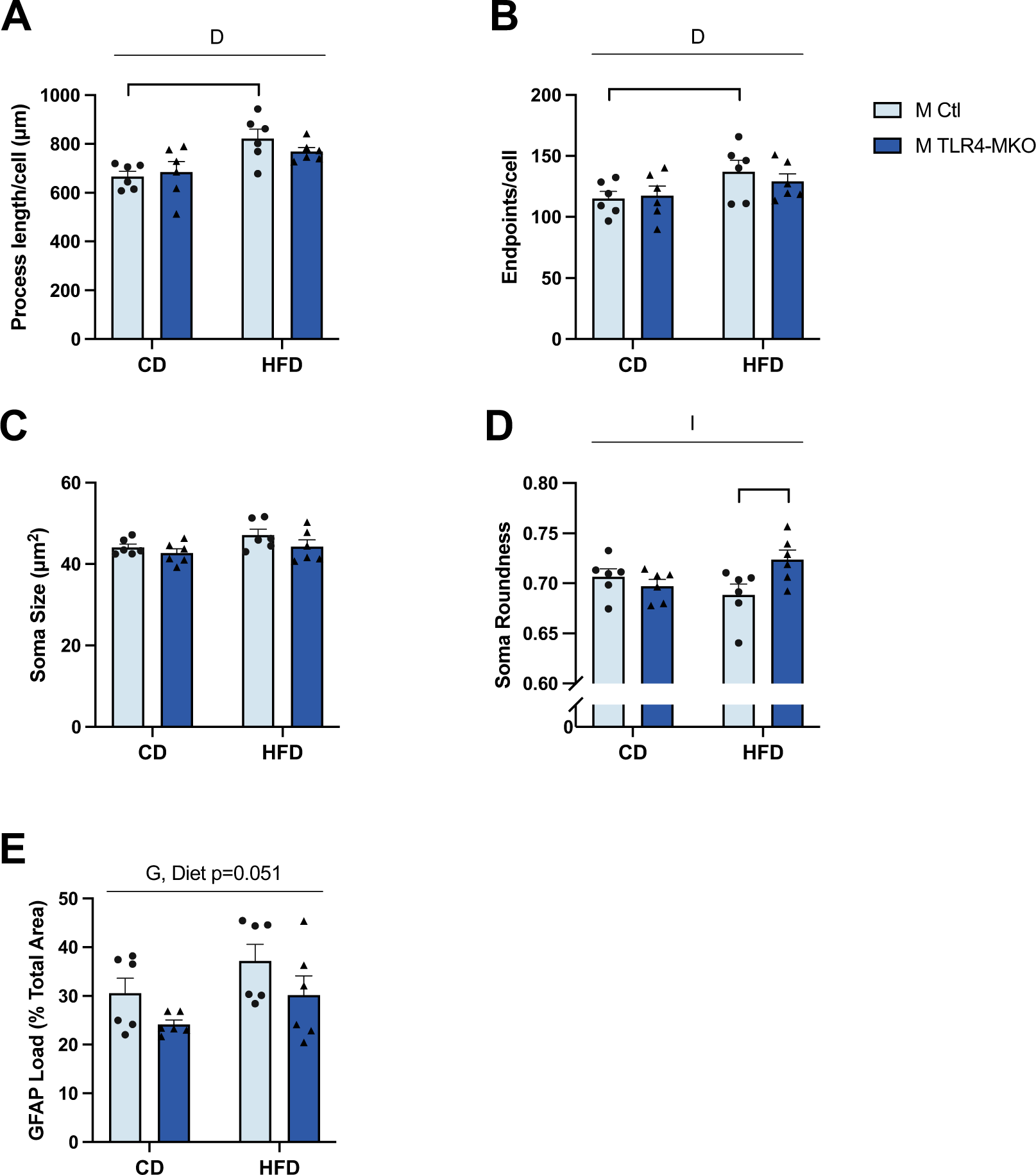
Effects of diet and microglial TLR4 deletion on microglia morphology and astrocyte immunoreactivity in the hippocampus of male mice. Images of Iba1 immunoreactivity from male control (Ctl, light blue bars) and TLR4-MKO (dark blue bars) mice following exposure to control (CD) or high-fat (HFD) diets were assessed for (A) average process length of Iba1+ cells, (B) average process endpoints per cell of Iba1+ cells, (C) soma size and (D) soma roundness of Iba1+ cells. (E) Immunoreactive loads of the astrocyte marker glial fibrillary acidic protein (GFAP) were also quantified. Data are presented as mean (+SEM) values and include individual values (filled circles for Ctl, filled triangles for TLR4-MKO); n = 6/group. Statistically significant main effects are denoted by D (diet), G (genotype), and I (diet and genotype interaction); significant between group comparisons of interest are identified with brackets and significance levels are denoted by * *p* < 0.05.

In female mice, we observed no significant effects of genotype and diet on markers of microglia morphology and astrocyte reactivity in the hippocampus (Figure 15).

**Figure 15.**
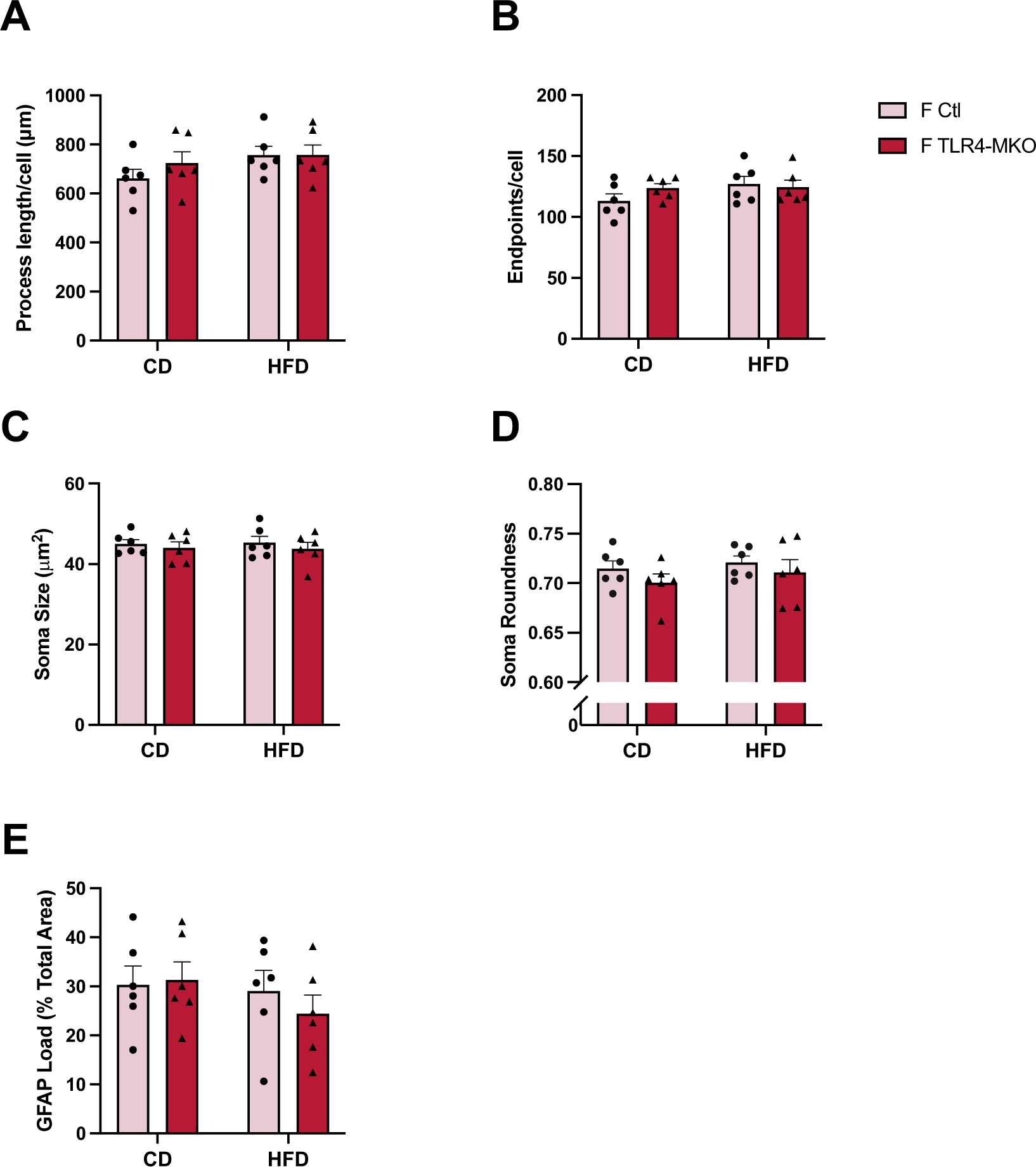
Effects of diet and microglial TLR4 deletion on microglia morphology and astrocyte immunoreactivity in the hippocampus of female mice. Images of Iba1 immunoreactivity from female control (Ctl, pink bars) and TLR4-MKO (red bars) mice following exposure to control (CD) or high-fat (HFD) diets were assessed for (A) average process length of Iba1+ cells, (B) average process endpoints per cell of Iba1+ cells, (C) soma size and (D) soma roundness of Iba1+ cells. (E) Immunoreactive loads of the astrocyte marker glial fibrillary acidic protein (GFAP) were also quantified. Data are presented as mean (+SEM) values and include individual values (filled circles for Ctl, filled triangles for TLR4-MKO); n = 6/group. No statistically significant main or interactive effects were observed.

### 3.9 Effect of HFD and microglial TLR4 on cognitive performance

Prior work with TLR4 knockouts demonstrated that TLR4 has important functions in brain development and behavior. For example, mice with constitutive knockout of TLR4 have impaired motor coordination and the deficiency is associated with a reduction in the thickness of the molecular layer of the cerebellum^75^. Moreover, TLR4 antagonist treated mice score higher on measures of anxiety^76^. Therefore, we determined if there was a behavioral phenotype associated with our induced microglial TLR4 deletion one week prior to dietary treatment. There were no significant genotype differences in any of the behavioral outcomes, including exploratory activity in open field (Figure S3A-C) and anxiety-like behavior in elevated plus maze (Figure S3D-E).

To investigate whether macrophage/microglial TLR4 deletion modulates HFD-associated cognitive outcomes, we examined behavior on the Barnes maze, a task of hippocampal-dependent spatial learning and memory. We first evaluated learning in males by measuring the latency to reach the escape box during training trials. Male mice across all groups showed significant learning with increased training, as indicated by a significant main effect of time (*F* _(3, 147)_ = 67.4, *p* < 0.001; Figure 16A) with shorter latencies to reach the escape box on days 2-4 than on day 1 of training (*p* < 0.05). Neither diet nor genotype significantly affected rates of learning. In the probe trial, we found a significant interaction between diet and genotype on primary latency in male mice (*F* _(1, 49)_ = 4.8, *p* = 0.033; Figure 16B) with a statistically nonsignificant trend toward longer latency in Ctl mice on HFD compared with Ctl mice fed on CD (*p* = 0.13) and a nonsignificant trend of shorter latency in TLR4-MKO mice versus Ctl mice under the HFD condition (*p* = 0.11). We also found a significant main effect of genotype on percent errors (*F* _(1, 49)_ = 9.3, *p* = 0.004; Figure 16C) in which TLR4-MKO mice showed fewer errors when selecting the correct hole than Ctl mice.

**Figure 16.**
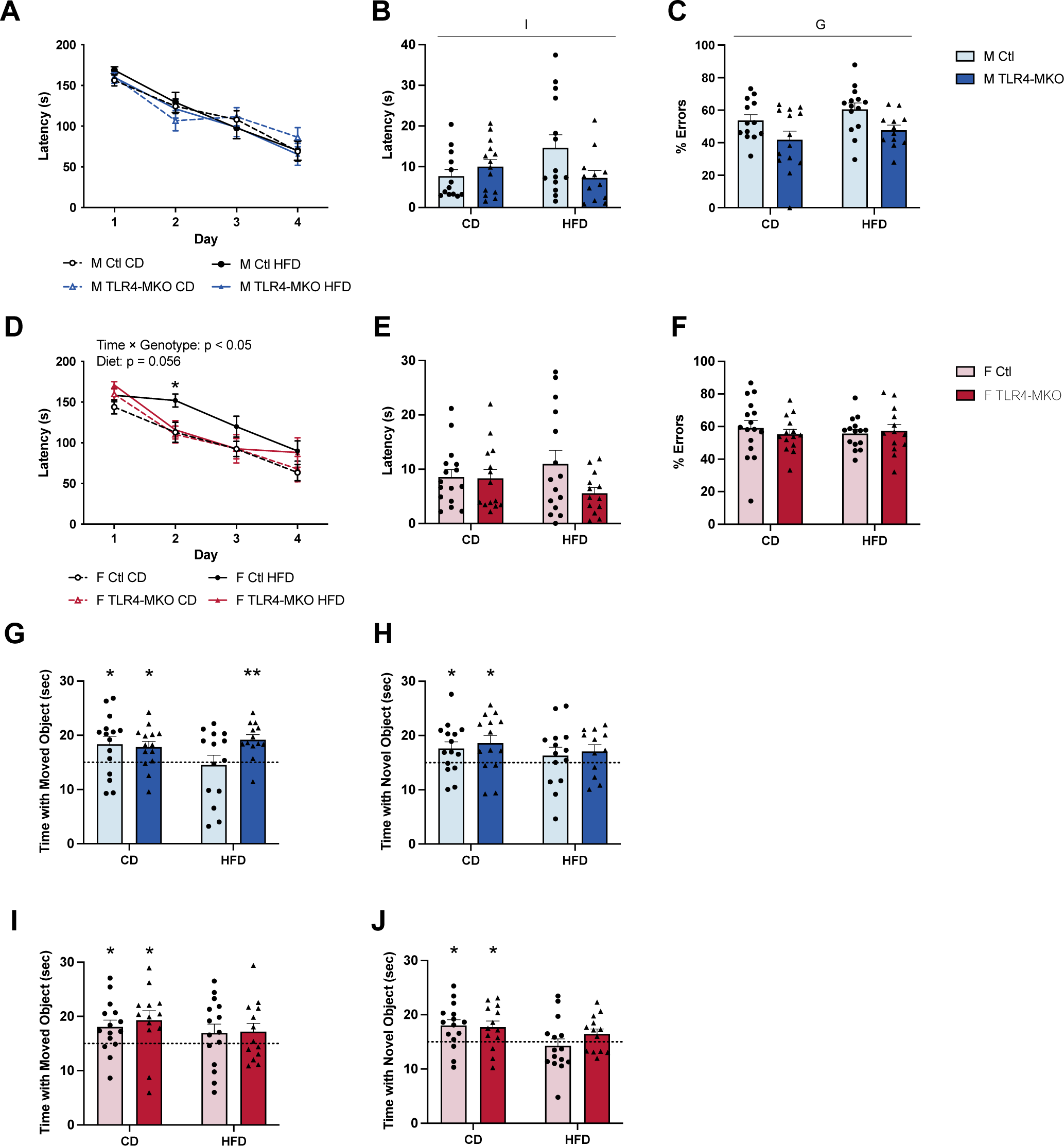
Effects of diet and microglial TLR4 deletion on cognitive performance in male and female mice. (A-C) Male control (Ctl, light blue bars) and TLR4-MKO (dark blue bars) and (D-F) female control (Ctl, pink bars) and TLR4-MKO (red bars) were behaviorally assessed following exposure to control (CD) or high-fat (HFD) diets for spatial learning and memory using the Barnes maze: (A, D) average escape latency during the 4 days of training, (B, E) prime latency in probe trial, and (C, F) percent errors made during the probe trial. The mice were also assessed on 30-second novel object placement (G,I) and recognition (H, J) tasks; dotted lines in indicate exploration at chance level (15 sec). Data show times spent with novel objects. All data are presented as mean (+SEM) values and include individual values (filled circles for Ctl, filled triangles for TLR4-MKO); n = 12-16/group. Statistically significant main effects are indicated by G (genotype) and I (diet and genotype interaction) with significance denoted by * *p* < 0.05, ** *p* < 0.01.

Female mice also showed significant learning during training trials, as there was a significant main effect of time (*F* _(2.440, 131.7)_ = 75.8, *p* < 0.001; Figure 16D). Additionally, we found a significant interaction between time and genotype (*F* _(2.440, 131.7)_ = 3.6, *p* = 0.022) and a non-significant tread toward an effect of diet (*p* = 0.056). Between group comparisons revealed that Ctl mice on HFD were slower at locating the escape box than CD mice specifically on day 2 of the training (*p* = 0.01). There was no main effect of diet nor an interaction between genotype and diet on probe latency or errors in female mice (Figure 16E-F).

We also examined cognitive performance using novel object placement (OP) and object recognition (OR) tasks. For male mice, obesity impaired behavioral performance such that Ctl mice on HFD had impaired performance compared with CD diet in both OP and OR tasks (Figure 16G-H). TLR4-MKO mice on HFD performed better than control HFD mice as they spent significantly more time than chance with the object in the novel location (*p* < 0.05), suggesting absence of microglial TLR4 improves cognition in obese male mice. For female mice, HFD was associated with impaired performance in OP and OR tasks regardless of genotype (Figure 16I-J).

**Figure S3.**
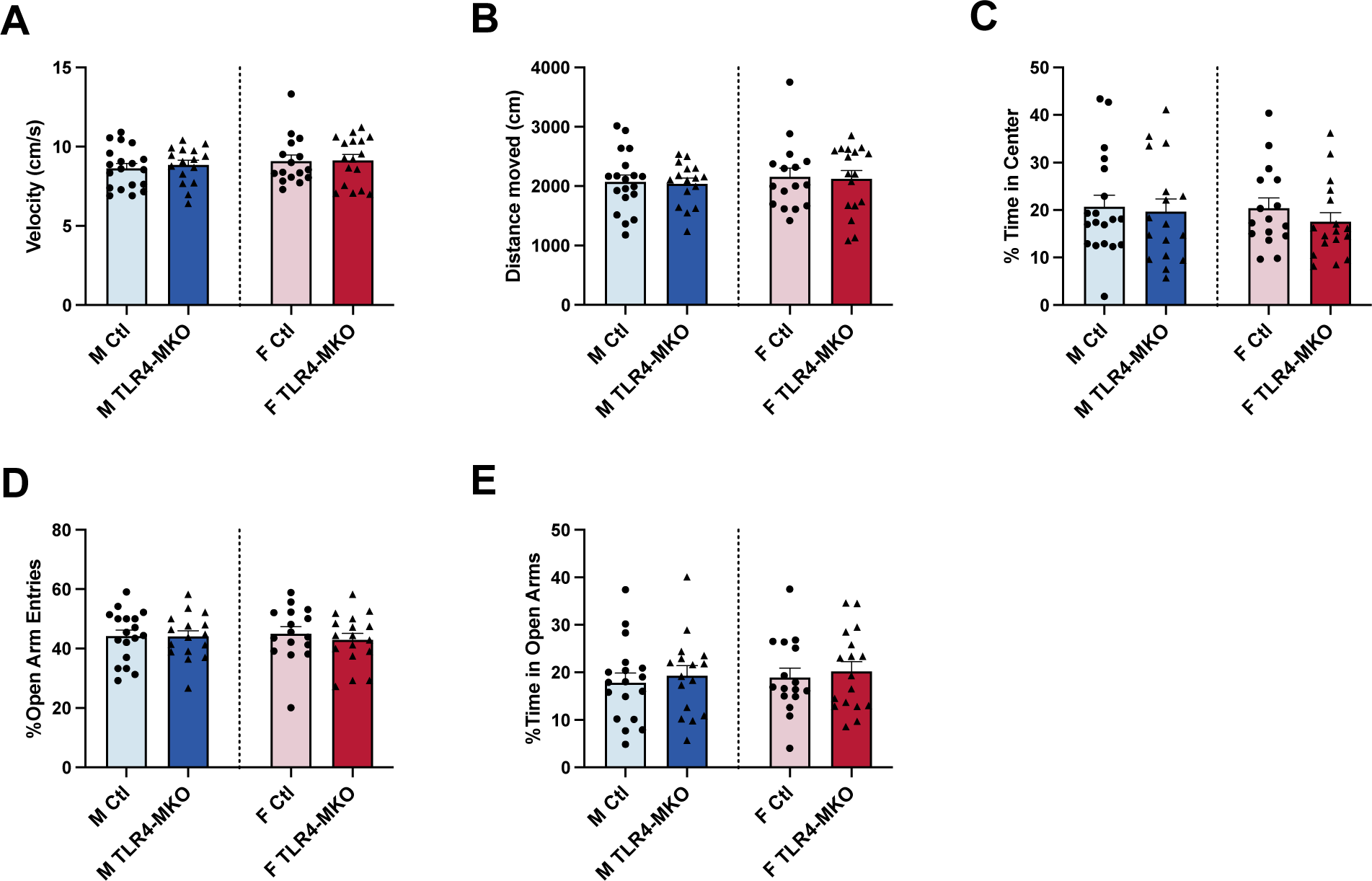
Effects of microglial TLR4 deletion on exploration and anxiety-like behaviors in male and female mice. Male control (Ctl, light blue bars) and TLR4-MKO (dark blue bars) and female control (Ctl, pink bars) and TLR4-MKO (red bars) were behaviorally assessed for (A-C) explorative and anxiety-like behaviors were examined in the open field: (A) average velocity, (B) total distance traveled, and (C) the amount of time spent in the center field. Anxiety-like behavior was assessed in the elevated plus maze (D,E): (D) percentage of arm entries into the open arm, and (E) the amount of time spent in the open arm. Data are presented as mean (+SEM) values and include individual values (filled circles for Ctl, filled triangles for TLR4-MKO); n = 16-19/group. No statistically significant main or interactive effects were observed.

### 3.10 Microglial TLR4 deletion improved hippocampal neurogenesis in obese male mice

Finally, we assessed neurogenesis, a process positively associated with hippocampal-dependent behavioral performance^77^, and known to be inversely impacted by obesity^78,79^. Numbers of new neurons in the dentate gyrus region of the hippocampus were measured by quantifying cells positive for doublecortin (DCX+) immunolabeling. In male mice, we found a significant main effect of diet (*F* _(1, 20)_ = 6.1, *p* = 0.023) and an interaction between diet and genotype (*F* _(1, 20)_ = 5.8, p = 0.026) on DCX+ cell numbers (Figure 17E) whereby HFD was significantly associated with impaired neurogenesis only in Ctl mice. We also assessed the maturation of DCX+ cells by examining their morphology. No main effects or interactions were found on the proportion of immature (type 1) and less immature (type 2) DCX+ cells. However, HFD-fed mice had significantly less mature cells (type 3) compared with mice on CD (*F* _(1, 20)_ = 6.6, *p* = 0.018; Figure 17F).

**Figure 17.**
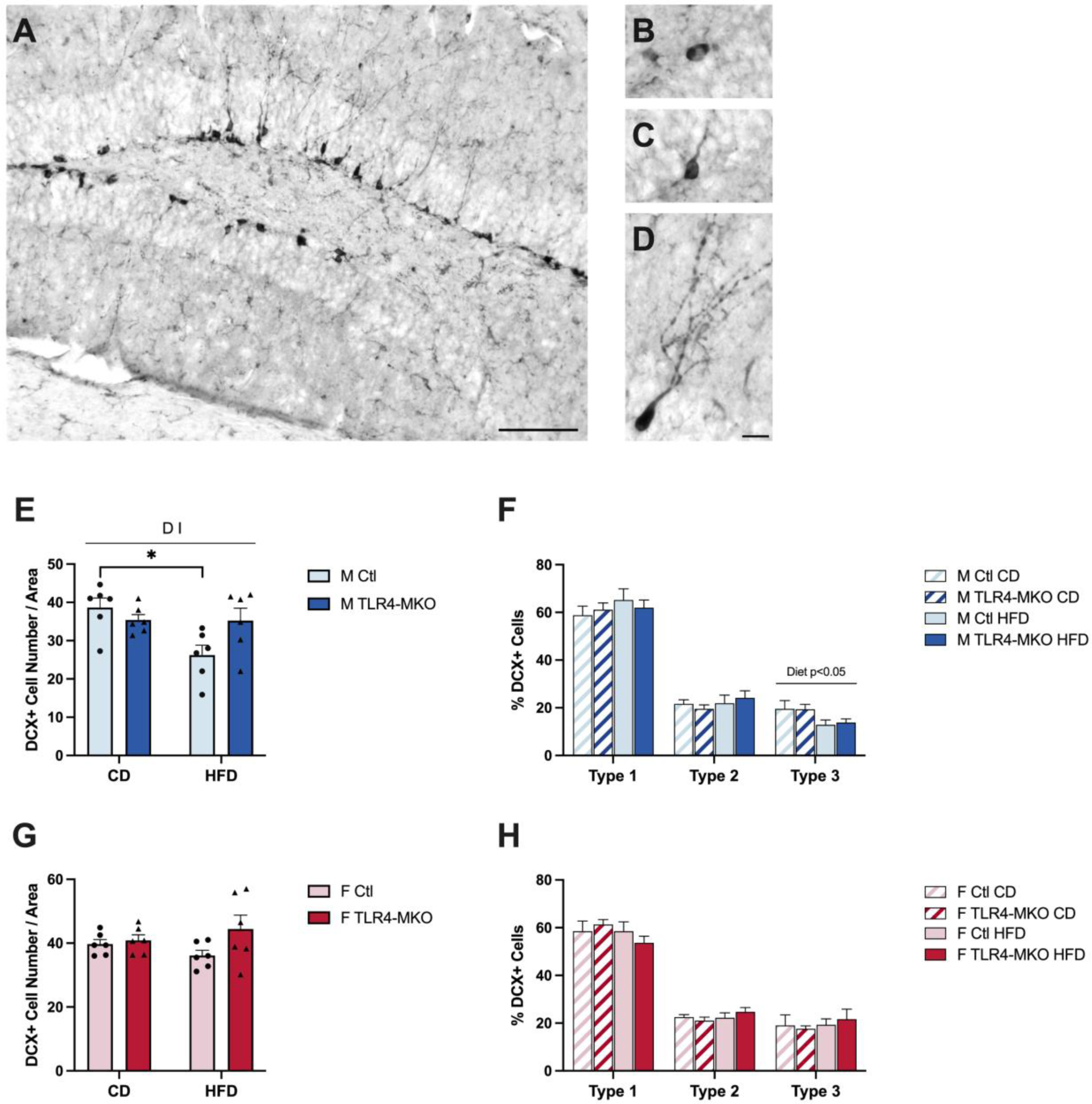
Effects of diet and microglial TLR4 deletion on hippocampal neurogenesis in male and female mice. (A) Representative image of doublecortin (DCX) immunoreactivity, a marker of newly differentiated neurons, in the dentate gyrus of the hippocampus; scale bar = 100 μm. Representative images of morphological categories of DCX-labeled cells as Type 1 (B), Type 2 (C), and Type 3 (D); scale bar = 15 μm. Images of DCX immunoreactivity from (E) male control (Ctl, light blue bars) and TLR4-MKO (dark blue bars) mice and (G) female control (Ctl, pink bars) and TLR4-MKO (red bars) mice following exposure to control (CD) or high-fat (HFD) diets were quantified for (E, G) numbers of DCX-labeled cells per unit area and (F, H) the relative proportion of DCX-labeled cells in the three morphological subtypes. Data are presented as mean (+SEM) values and, where appropriate, include individual values (filled circles for Ctl, filled triangles for TLR4-MKO); n = 6/group. Statistically significant main effects are denoted by D (diet) and I (diet and genotype interaction); significant between group comparisons of interest are identified with brackets and significance levels are denoted by * *p* < 0.05.

Parallel examination of neurogenesis in female mice showed no significant differences across groups in the numbers or relative maturity of DCX+ cells (Figure 17G-H).

## 4. Discussion

In this study, we examined the effects of microglial TLR4 deletion on systemic and neural changes induced by 16-week HFD exposure in male and female mice. The data demonstrate that TLR4-MKO mice were protected against numerous HFD-associated impairments, including metabolic imbalance, peripheral inflammation, cognitive impairment and altered neurogenesis. Interestingly, the beneficial effects of microglial TLR4 deletion were sexually dimorphic. Specifically, we found that although HFD induced very similar metabolic outcomes in male Ctl and TLR4-MKO mice, microglial TLR4 deletion yielded protection from HFD-induced adipose inflammation, neuroinflammation, cognitive impairment and diminished neurogenesis. Conversely, female TLR4-MKO mice exhibited lower body weight gains and improvements in some metabolic functions relative to Ctl mice in the context of obesogenic diet, yet showed limited evidence of improved neural function. These findings provide insights into the critical role of TLR4 in modulating HFD-induced metabolic disorder and inflammation in a sex-dependent manner.

Prior studies have established strong connections among obesity, CNS inflammation, and metabolic dysfunctions, with obesogenic diets known to yield numerous metabolic disturbances including insulin resistance and increased adiposity. The role of neuroinflammation, mediated largely by astrocytes and microglia, in exacerbating systemic metabolic imbalances is increasingly recognized^80^. One well supported idea is that inflammatory signaling in the brain contributes to diet-induced metabolic disorders and drives peripheral inflammation^33,81–84^. Consistent with this position, pharmacological inhibition of IKKβ/NF-kB signaling in the CNS^85^, genetic knockout of IKKβ in the neurons^86^ and knockout of Myd88 in the CNS^87^ all reduced HFD-induced weight gain and insulin resistance. Additionally, intracerebroventricular (ICV) infusion of the pro-inflammatory cytokine TNFα increased insulin secretion and impaired insulin sensitivity in peripheral tissues, whereas blocking hypothalamic TNFα signaling prevented HFD-induced weight gain and insulin resistance^88^. Similarly, ICV administration of IL-4 exacerbated the hypothalamic inflammation and systemic metabolic outcomes of HFD, effects that were abolished by pre-treating the mice with an IKKβ inhibitor^89^.

Microglia have been implicated as key mediators of the mechanisms that link CNS inflammation and systemic metabolic function^82,90,91^. Microglia depletion was associated with reduced food intake induced by high dietary consumption of SFAs ^37,38^. Likewise, blocking microglia proliferation prevented HFD-induced weight gain, improved hypothalamic leptin sensitivity and decreased hypothalamic and peripheral inflammation^39^. Microglial-dependent TLR4 signaling contributes to HFD-induced neuroinflammation. For example, TLR4/MyD88 signaling was strongly activated in the hypothalamus after HFD exposure^28^. Moreover, TLR4 signaling is required for SFA-induced microglia activation^92^. Blocking TLR4 signaling attenuated HFD-induced expression of pro-inflammatory cytokines^28^. Our study advances this field by spotlighting microglial TLR4 signaling as a crucial determinant in the complex framework linking CNS inflammation and obesity-associated metabolic changes in both sexes. In male mice, deletion of microglial TLR4 did not significantly affect HFD-induced changes in most metabolic measures, including body weight, food intake, adiposity, and glucose tolerance. However, TLR4-MKO mice had significantly lower fasting insulin levels as well as reduced peripheral inflammation, as demonstrated by lower IL-β, IL-6, H2ab1 and CD74 levels in the visceral fat pad. There was a general pattern of modest, HFD-associated increases in plasma cytokines. However, parallel with our findings in adipose tissue, we observed lower levels of plasma IL-6 in TLR4-MKO male mice. This is in line with previous findings that circulating IL-6 is largely contributed by adipose tissue^93^. Conversely, in female mice, the deletion of microglial TLR4 yielded more profound effects. Notably, it was associated with significant lowering in body weight, improvements in metabolic function and reduction in systemic inflammation. These results are consistent with previous findings highlighting the pivotal role of microglial TLR4 signaling pathway in modulating neuronal activity and feeding behavior within the ARC^94^.

IL-6 is a recognized immune modulator that is known to exert significant influence on various aspects of CNS function^95^. In contrast to the lower peripheral IL-6 levels observed in TLR4-MKO mice, our study revealed a noteworthy association between the deletion of microglial TLR4 and heightened mRNA expression of IL-6 within the hippocampus and hypothalamus. Prior studies have shown that IL-6 signaling in the CNS plays an important role in regulating food intake and metabolic functions. For example, central administration of IL-6 significantly reduced food intake, improved glucose tolerance, and increased brown adipose tissue thermogenesis in lean as well as obese mice^96,97^ likely by suppressing hypothalamic IKKβ activation and ER stress^98^. On the other hand, inhibition of IL-6 expression in the lateral parabrachial nucleus increased food intake, accompanied with increased body weight and adiposity^97^. CNS IL-6 is also involved in the control of neuronal functions such as neurogenesis^99^, synaptic plasticity^100^ and learning and memory^101^. Because of the divergent pattern of IL-6 levels in the plasma and brain, our findings suggest different functions of IL-6 signaling in the CNS versus peripheral tissues. Indeed, IL-6 signaling promotes obesity-induced adipogenesis and macrophage accumulation in the adipose tissue^102^. Moreover, chronic subcutaneous infusion of IL-6 impaired hepatic insulin signaling and induced insulin resistance^103^. Interestingly, a recent paper showed that IL-6 secreted from adipocytes promoted, while IL-6 expressed by myeloid cells (including macrophages and microglia) suppressed, HFD-induced metabolic impairment and adipose tissue inflammation^104^. These data suggest that the outcomes of IL-6 signaling are cell type dependent, a possibility that would reconcile the apparent discrepancy of anti- versus pro-inflammatory characteristics of IL-6 in the CNS relative to the periphery. In the CNS, IL-6 is expressed by many cell types including neurons, microglia, astrocytes and endothelial cells^95^. Though the exact cell contribution of observed increases in IL-6 expression remains to be established, it is possible that elevated IL-6 signaling contributes to improved metabolic function in TLR4-MKO mice.

This study significantly advances our understanding of the important roles of microglial TLR4 signaling in the multiple adverse CNS consequences of obesity. We began by examining its contribution to activated phenotypes in both astrocytes and microglia, which in turn may impact cognition. Our study supports the hypothesis that microglial TLR4 signaling modulates HFD-induced microgliosis and astroglial activation. First, we observed that HFD increased expression of microglial specific marker Tmem119 and Trem2 in male mice, with a general pattern that trended towards a significant increase only in Ctl animals. The only exception was that no diet effect was found on Trem2 levels in the hypothalamus, which might be due to microglia heterogeneity and a much lower Trem2 expression in the hypothalamus than other brain regions^71^. A transcriptome study using RNA-seq has demonstrated downregulation of Tmem119 and Trem2 in isolated microglia after HFD exposure^73^. However, different expression pattern associated with microglial activation and inflammation was observed when examining mRNA changes on the level of whole tissue. HFD is associated with elevated expression of Trem2 in the hippocampus^105^. In addition, expression of Tmem119 and Trem2 was upregulated in the hippocampus and was further increased by HFD feeding in Alzheimer’s disease mouse models^106,107^. Though the exact function and gene expression pattern of Tmem119 and Trem2 under pathological conditions need to be further elucidated, HFD-associated upregulation of these genes observed in our study may reflect an increase in microglia cell number rather than altered activation state. Second, we found attenuated HFD-induced microglial activation in TLR4-MKO mice in both sexes, demonstrated by partial reductions in markers of microglia activation (H2Ab1 and CD74), as well as changes in microglia morphology. Third, TLR4-MKO mice had significant less astrocyte activation in the hippocampus and hypothalamus in male mice and in the hypothalamus in female mice, suggesting microglia-astrocyte interactions in modulating neuroinflammation.

Beyond neuroinflammation, our findings inform on the role of microglial TLR4 signaling in diet-induced cognitive deficits. Several studies have underscored the significant association between microglial activation and cognitive function. For example, rats on HFD showed significant cognitive impairments along with altered microglial morphology^108^. Inhibition of microglial activation by minocycline treatment or blocking microglial phagocytosis did not affect HFD-induced body weight gain, but did improve performance in hippocampal-dependent behavioral tasks^44^. Moreover, another study showed that switching from a HFD back to a low-fat diet reduced microglial activation and attenuated the synaptic plasticity in obese mice without complete reversal of body weight^109^. These findings suggest a stronger relationship between cognitive function and microglial activation than with adiposity. Our study corroborates this link, as microglial TLR4 deletion is shown to enhance certain behavioral measures, offering valuable insights into its role in mitigating cognitive impairments associated with obesity.

Furthermore, we focused on relationships between TLR4 signaling and neurogenesis. Activation of TLR4, along with associated pathways of NF-κB activation, have been demonstrated to exert negative effects on proliferation of neural progenitor cells and neurogenesis^110,111^. Moreover, microglia have an important role in regulating generation of new neurons both under physiological condition and stress^112–114^. When examining neurogenesis in the hippocampus, we found that DCX^+^ cell numbers in the dentate gyrus were not significantly affected by HFD in female mice. In contrast, HFD impaired hippocampal neurogenesis in male mice, an effect attenuated by TLR4 deletion.

We also evaluated sex differences in outcomes associated with HFD and microglial TLR4 deletion. We found that microglial TLR4 deletion did not ameliorate the majority of the metabolic dysregulation induced by HFD feeding in male mice. In contrast, female obese mice with microglial TLR4 deficiency had improved metabolic function which was likely affected by lower caloric intake. Many studies have found that female sex is generally protected from the development of obesity and HFD-induced metabolic alterations in young adult rodents^117–119^, however with prolonged dietary treatment and or increased age female mice showed similar levels of body weight gain, insulin resistance and adipose inflammation as males^120,121^. Although the exact mechanisms remain unknown, hypothalamic neural circuits controlling energy balance have been proposed to attribute to the delayed response to HFD in female mice. Activation of the POMC neurons suppresses food intake and increases energy expenditure^122,123^. Female mice have higher expression of POMC in the hypothalamus^124–126^, and their POMC neurons exhibit higher neural activity^125^. The protection against obesity in female mice might also be due to inherent sex differences in microglial function and characteristics. During early development, microglial numbers, morphology, and activation states show notable differences between male and female brains^127^. In terms of function, sex-related differences in microglia are evident in phagocytosis and antigen presentation, especially in the context of neuroinflammatory diseases like Alzheimer’s disease. Research has shown that microglia from female APP/PS1 mice, tend to be more glycolytic and less phagocytic compared to male mice^128^. At the transcriptomic level, male sex is significantly associated with upregulation of NF-κB and inflammatory processes^129^. Female mice with CX3CR1 deficiency displayed a ‘masculinized’ response to HFD, where they gained significantly more weight and displayed higher microglial activation and inflammation than their control littermates^130^. Moreover, TLR4 has recently been implicated in these sex differences. Studies suggest a significant role for TLR4 in modulating microglial functions differently in males and females^131,132^. Our data also show that HFD affected gene expression and glial activation in a sex-dependent manner. In male mice, HFD altered outcomes in both hippocampus and hypothalamus, whereas in female mice HFD did not affect any of the neuronal outcomes in the hippocampus. These data suggest that neuroinflammation induced by 16-week HFD treatment is not generalized to the whole brain in females. Indeed, inflammation induced by HFD develops sooner in the hypothalamus than other brain regions. In male rodents, expression of inflammatory cytokines in the hypothalamus was elevated within 24 hours of HFD consumption, while changes were not observed in the hippocampus after up to 12 weeks of HFD^36,109,133^.

Even though female sex is more resistant to diet-induced metabolic impairment and inflammation, they are not protected from obesity-related cognitive impairment^134,135^. We found sex differences on some of the behavioral outcomes. HFD significantly impaired learning behavior during Barnes maze training phase only in female Ctl mice. Additionally, microglial TLR4 deletion improved performance in the object placement test in male but not female mice. Previous studies have suggested sex differences in hippocampal-dependent spatial navigation and learning tasks in rodent models, with some studies suggesting a male bias ^136–138^. Similarly, sex differences on spatial learning and memory performance have been widely reported in humans, with evidence of males outperforming females on spatial tests ^139–142^. Thus, our observed differences in behavioral measures may suggest inherent sexually dimorphic responses in spatial tasks. Additional research is needed to further define how sex interacts with microglial inflammation in regulating DIO as well as the underlying mechanisms of these interactions.

There are some limitations to the study that may impact observed metabolic and sex-dependent effects. In our mouse model, tamoxifen triggered TLR4 deletion specifically in Cx3CR1-expressing cells. In the brain, Cx3CR1 expression is limited to microglia^54^. At adult ages, Kupffer cells in the liver, peritoneal, splenic and lung macrophages do not express Cx3CR1^40^. Although circulating monocytes and a subpopulation of macrophages residing in the intestine have been reported to express Cx3CR1 during adulthood, they are short-lived with half-lives of 2 days and 3 weeks, respectively^40,41^. In lean mice, only a small population of macrophages in the adipose tissue (∼6%) is positive for CX3CR1^42^. After 8 weeks of HFD feeding, the CX3CR1^+^ cell population was largely expanded, but was mostly contributed by infiltrating macrophage^42^. In our study, tamoxifen treatment did not reduce TLR4 expression in the adipose tissue in TLR4-MKO female mice, and unexpectedly resulted in non-significantly increased TLR4 expression in males. The latter might be a result of compensatory upregulation of TLR4 in other cell types, such as cells in the stromal vascular fraction^23^, after transient deletion of TLR4 in circulating monocytes/macrophages. Nonetheless, we cannot rule out potential effects of peripheral tissue-resident macrophages on observed effects of TLR4 deletion on HFD-induced metabolic function and systemic inflammation.

## 5. Conclusion

In summary, our data show sex-dependent protective effects of microglial TLR4 signaling against adverse HFD-induced outcomes. In male mice, deletion of microglial TLR4 improved insulin resistance, inhibited adipose inflammation and increased neurogenesis. The reduced adiposity and an overall improvement of metabolic functions in female TLR4-MKO mice might have resulted from lowered food intake, which suggested links between microglial TLR4 signaling and hypothalamic neuronal circuits in regulating energy balance. Obese mice with microglial TLR4 deletion exhibited significant protection against diet-induced cognitive impairments and glial activation in both male and female mice. These findings add to the growing literature suggesting an important role for microglial inflammation in mediating obesity-related outcomes. Clinical approaches targeting microglial TLR4 signaling may be useful as neuroinflammation and obesity therapeutics.

## Acknowledgements

This work was supported in part by a grants from the NIA (AG055367). The authors thank Dr. Amy Christensen and Ms. Cassandra McGill for helpful comments on the manuscript.

